# Tau pSer396 and pSer404 Define Distinct Epitope Regions Linked to Different Antibody Functions

**DOI:** 10.64898/2026.04.16.716390

**Authors:** Ruimin Pan, Erin E. Congdon, Jessica E. Chukwu, Christina C. Luo, Einar M. Sigurdsson, Xiang-Peng Kong

**Affiliations:** Department of Biochemistry and Molecular Pharmacology, New York University Grossman School of Medicine, New York NY 10016, USA; Department of Neuroscience, Institute for Translational Neuroscience, New York University Grossman School of Medicine, New York NY 10016, USA; Department of Psychiatry, New York University Grossman School of Medicine, New York NY 10016, USA

## Abstract

Hyperphosphorylated tau is a central pathological feature of Alzheimer’s disease and related tauopathies, and antibodies targeting the pSer396/pSer404 epitope region represent a promising therapeutic strategy. However, direct comparisons of pSer396- and pSer404-selective antibodies and the impact of humanization on their functional properties remain limited. We generated two new monoclonal antibodies (mAbs), 9E (pSer404-specific) and G10 (pSer396-specific), and evaluated them alongside 4E6 (pSer404) and PHF-1 (pSer396) in murine and partially humanized chimeric formats. Antibodies were assessed in mixed cortical cultures using extracellular (PHF + Ab) and intracellular (PHF → Ab) paradigms. Efficacy in preventing tau-induced toxicity and seeding differed substantially among antibodies and was variably altered by chimerization, despite identical variable regions. Antibodies targeting pSer404 were more effective than those targeting pSer396, and antibodies that preferentially bound soluble pathological tau species in competition ELISA were consistently more efficacious, whereas neuronal uptake was comparable across variants. To define structural determinants of phospho-epitope recognition, we determined the crystal structures of the Fab regions of 9E, G10, and PHF-1, and additionally solved the co-crystal structure of Fab PHF-1 in complex with a pSer396 tau peptide at 2.55 Å resolution. The PHF-1 complex reveals a heavy-chain-dominant binding mode in which pSer396 is anchored within an electropositive pocket and Tyr394 adopts a flipped conformation that stabilizes a β-strand-like motif, consistent with a phosphorylation-dependent conformational switch. These findings demonstrate that epitope selectivity, aggregate preference, structural binding mode, and Fc context collectively govern antibody efficacy, and that humanization can substantially alter therapeutic properties.

## Introduction

Under normal conditions, the tau protein exists as a natively unfolded monomer that is stable and soluble, even at high temperatures, providing cytoskeletal stability to neuronal axons and acting as a regulator of transport (for a recent review see [1]). However, in tauopathies, the protein is heavily post-translationally modified, including hyperphosphorylated, leading to the development of toxic assemblies [2, 3]. Tauopathies, of which Alzheimer’s disease (AD) is the most common, present with unique patterns of tau phosphorylation and aggregation that can change with disease stage [4–10]. Hyperphosphorylation reduces both tau binding to microtubules and promotes misfolding, which can lead to self-assembly, forming first soluble oligomers, followed by larger insoluble filaments that make up the neurofibrillary tangles. These oligomers are the primary toxic species and play a key role in tau seeding, neuronal toxicity, and behavioral impairments [11–13]. Because of the key role that phosphorylation plays in the development of tauopathies, considerable effort has been on developing approaches to reduce tau phosphorylation, or promote the clearance of hyperphosphorylated tau, including with tau immunotherapy.

Since the first reports from our group [14–17], tau immunotherapy has advanced with both active and passive approaches in clinical trials, and remains a promising avenue for the development of therapeutics (for recent reviews see [18–20]). Although multiple tau antibodies (Abs) have reached clinical trials, none have yet been approved as a treatment for AD or other tauopathies. There are various challenges to designing a successful Ab and bringing it to human patients, including the choice of subclass and epitope, as well as the necessary process of humanization. Regarding epitope, various ones have been targeted, including phospho-epitopes, misfolded or oligomeric tau, or non-phosphorylated linear regions of tau. Of these, one of the most frequently selected is the phospho (p)Ser396/404 site [19, 21], which we focused on in our initial reports. The prevalence of this epitope in all tauopathies has made it of special interest.

pSer396 and pSer404 are present in both intra- and extracellular pathological tau. These closely aligned phospho-residues are elevated in the brain of all tauopathy patients, and in the cerebrospinal fluid (CSF) of AD patients [3, 4, 22–28]. Interestingly, the pSer396/404 site lies adjacent to the filament core in the “fuzzy coat” region, where tau adopts multiple conformations [29]. Phosphorylation at this site may lead to early pathological changes in tau by disrupting proper tau localization, initiating tau oligomerization, and facilitating tau accumulation. Both these p-sites reduce microtubule binding, inhibit tau’s ability to promote tubulin assembly and promote conformational changes in tau [30–34]. Additionally, pSer396/404 epitopes can form β-sheet structures, which may play a role in the seeding core of tau oligomers [35, 36]. An additional interesting feature is that in vitro phosphorylation at Ser396 requires prior phosphorylation of Ser404 [37, 38].

While typically considered parts of the same epitope, Abs can be made that are selective for one phospho-residue or the other. C10.2 and its humanized version are highly selective for pSer396 [39, 40]. The murine C10.2 has been shown to reduce tau seeding when tau was preincubated or immunodepleted with the Ab, and to reduce pSer396 in the CSF of transgenic mice [28, 39]. In cultured glia, C10.2 promoted tau phagocytosis in an Fc-dependent manner [41]. The Ab has entered clinical trials under the name Lu AF87908. PHF-1, another pSer396 selective Ab cleared pathological tau and improved function in a tauopathy mouse model [16, 17, 42]. Ittner et al. showed reduced tau pathology with another Ab targeting pSer404 [43]. We have previously published results using two pSer404 selective Abs 4E6 and 8B2. 4E6 is phospho-tau selective and does not bind to pSer396 [44]. It acutely improved cognition, reduced soluble phospho-tau, partially normalized neuronal calcium signaling in vivo, and prevented both toxicity and tau seeding intra- and extracellularly in culture [44–49]. 8B2 is also p-tau-selective and in ELISA showed binding to a pSer396 peptide only at high Ab concentrations [35]. Like 4E6, 8B2 was taken up into neurons and colocalized with tau following intravenous injection into tauopathy mice [35, 50]. It reduced soluble p-tau and gliosis and improved neuronal calcium signaling in these animals [50]. Another tau Ab 6B2 has comparable binding to the pSer396 and pSer404 epitopes as well as their non-p version in solid phase but has preference for the p-epitopes in solution. However, despite its potential utility as an imaging agent, it lacks efficacy to clear pathological tau, although it colocalizes with tau in brain neurons following its intravenous injection [51]. Its lack of efficacy, relative to 4E6 may relate to their different tau binding profiles, with 4E6 binding more readily to soluble tau species while 6B2 preferentially binds to larger insoluble aggregates [49]. Notably, 6B2’s higher affinity for solid phase tau may explain its lack of efficacy since its strong binding to tau in neuronal lysosomes may render these aggregates more compact and less accessible to lysosomal enzymes.

While Abs targeting one or the other p-site have shown efficacy in various models, a direct comparison of several pSer396 and pSer404 selective Abs has not been previously performed. Additionally, as stated above, one of the major challenges to creating disease modifying therapies is that the Abs need to be humanized which may change their properties. We have shown that human chimerization, and even changing the murine IgG subclass, can drastically alter the binding and efficacy of tau Abs despite the antigen binding region remaining the same [48, 52]. Thus, it is important to not only identify an epitope to target but also to examine how chimerization impacts Ab properties, including binding, neuronal uptake and ultimately efficacy.

To this end we generated murine and partially humanized chimera pairs of two novel Abs 9E and G10 targeting pSer404 and pSer396 respectively and compared them with mAbs 4E6 and PHF-1. Their efficacy in preventing tau-induced toxicity and seeding in mixed cortical cultures, as well as their cellular uptake and tau binding affinity were examined. To determine atomic level detail of the epitope regions, we also tried to determine crystal structures of the fragment of the antigen binding region (Fab) in complex with the tau peptide. For all the Abs, chimerization increased their isoelectric point, but the impact of the change differed. Consistent with our previous results [49, 52] murine 4E6 was effective both extra- and intracellularly and was negatively affected by chimerization. While murine PHF-1 was effective against extracellular tau, its chimera was not. On the other hand, chimerization did not affect the efficacy of 9E but appeared to be beneficial for G10. This further highlights the varying efficacy of Abs targeting this key pathological region of tau, with the pSer404 Abs examined here being more efficacious. Thus, it is important to thoroughly retest the efficacy of Abs following their humanization and before clinical trials.

## Materials and methods

### Animals

All animals were housed in IACUC approved facilities and experiments carried out using IACUC approved protocols. Pups from the homozygous JNPL3 mouse line (RRID:IMSR_TAC:2508) were used to generate the mixed cortical cultures. These animals express the human 0N4R tau isoform containing the P301L mutation in addition to wild type mouse tau [53].

### Generation of mouse monoclonal antibodies (mAbs)

Two groups of C5.26/BL6 mice at 2-3 months of age were immunized using DNA prime and protein boost immunization regimen. All mice were immunized three times with a DNA plasmid encoding full-length 2N4R tau cloned into the pJW4303 DNA expression vector and administered by skin tattooing [54, 55]. Then one group was protein boosted by a peptide of tau residues 391-407 (EIVYKpSPVVSGDTpSPRH) fused to an HIV-1 gp120 T1 epitope region (KQIINMWQEVGKAMYA) at the N-terminal and the other group was boosted by another peptide with tau residues 394-406 fused to a tetanus toxin epitope (GPSLFNNFTVSFWLRVPKVSASHLE) [56]. Mice were boosted 4 times intraperitoneally with 100 µg peptide formulated in incomplete Freund’s adjuvant (IFA) following the vendor’s protocol (Norcross, GA, USA). To test the immune responses, intermittent blood samples were collected 2 weeks after the last three boosts and tested by ELISA binding to the tau 386-408 (pSer396/pSer404) epitope peptide. The timeline of the immunization is shown in Figure 1. To generate the mAbs, the splenocytes of mouse number 2 in group 1 were used to generate hybridomas by a commercial vendor (ProSci Inc, Poway, CA). Hybridomas were selected for subcloning after exhibiting positive binding to ELISA plates that had been coated with 1 µg/mL peptide of tau 386-408 (pSer396/pSer404). Two hybridoma cell lines, 9E and G10, were generated. The mAb 4E6 was generated as previously described [44, 45]. The mouse PHF-1 mAb was generously provided by Peter Davies (Albert Einstein and Feinstein Institute, New York, NY).

**Figure 1.**
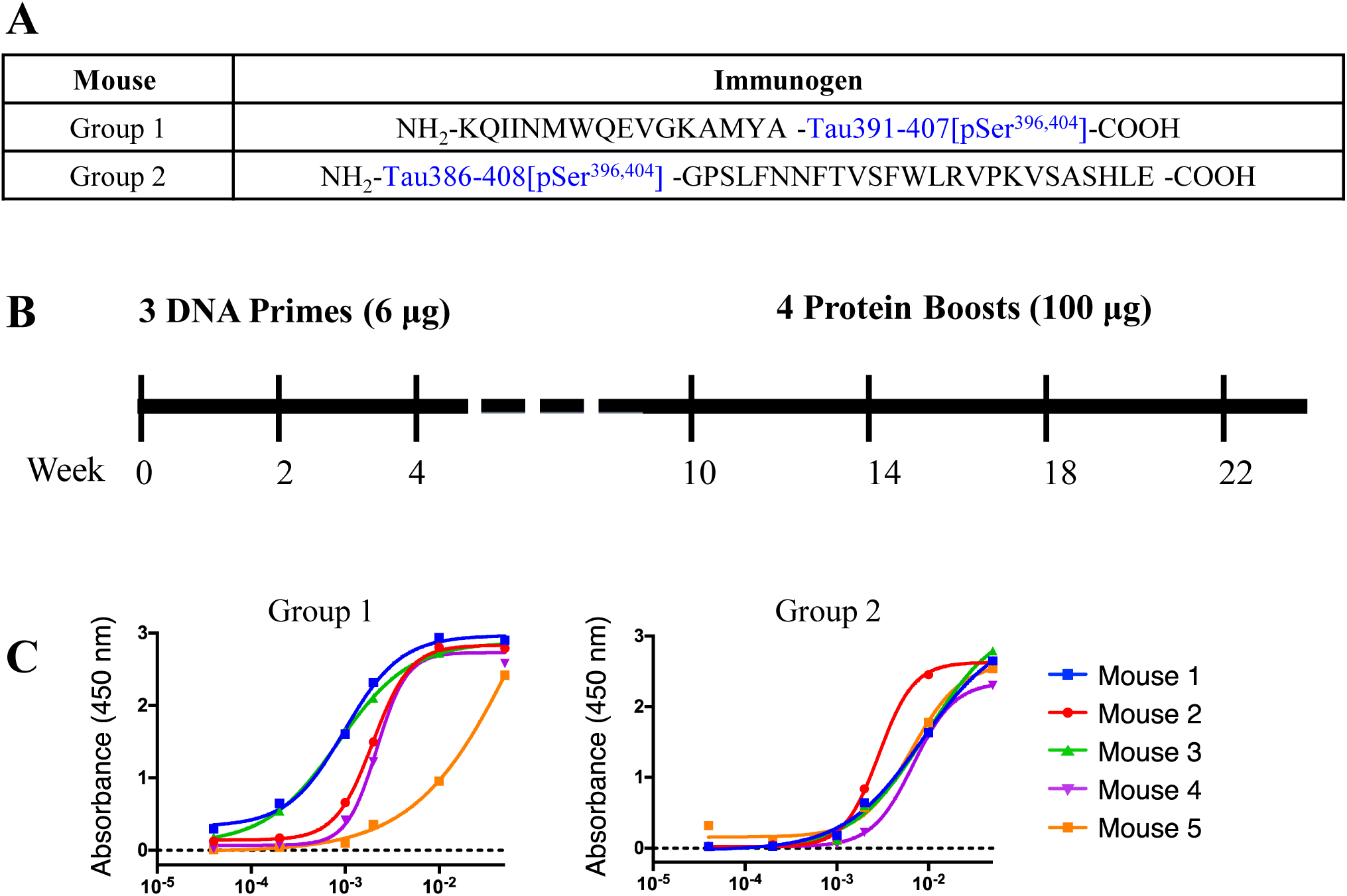
The immunization of mice with tau pSer396 and pSer404 epitope region. **A**. The two peptides used for the immunizations. **B**. The timeline of the immunization. Mice were immunized by the DNA prime-protein boost regimen. All mice were tattooed with DNA prime (6 µg) on the hind leg, then the mice were split into two groups and injected intraperitoneally with the two different peptides (100 µg). **C**. ELISA test of the immune responses at week 16 to the peptide _386_TDHGAEIVYKpSPVVSGDTpSPRHL_408_. Sera were serially diluted, and Ab titer was detected with an HRP-conjugated anti-IgG secondary antibody. Absorbance was measured at 450 nm. Mice in Group 1 showed stronger immune response. Therefore, mice in Group 1 were used to generate hybridoma. Both mAbs G10 and 9E were generated from Mouse 2 in Group 1.

### Antibody production and purification

The mAbs 9E and G10 were purified from the hybridoma cultures. The hybridoma supernatants were collected and purified by protein G column as per the manufacturer’s protocol (Cytiva, DE). Purified PHF-1 antibody was generously provided by Peter Davies (Albert Einstein College of Medicine and Long Island Jewish Medical Center, Litwin-Zucker Research Center at Feinstein Institutes for Medical Research). The mAb 4E6 was purified as previously described [44, 45]. The chimera G10, 9E, 4E6, and PHF-1 Abs were generated by cloning the heavy chain and light chain variable fragments into human IgG1 and IgGκ frames respectively [57]. The plasmids of the heavy chain and light chain were transfected into Freestyle^TM^ 293F cells (ThermoFisher Scientific, Waltham, MA). Cells were cultured for 5 days, after which the supernatants were purified by protein A/G column (Cytiva, DE). Their affinities to the p- and non-p Ser396/404 epitope region were determined by ELISA. For comparison, the recombinant chimera 4E6 was also produced in HEK293S GnTI^−/-^ cells (American Type Culture Collection, VA) that produce only high-mannose glycans, which are more homogeneous than complex glycans [58].

### Isoelectric focusing (IEF)

Each murine and chimeric Ab were added to IEF sample buffer and run on a precast 3–9 IEF gel (BioRad, Hercules, CA). The gel was stained overnight in a 20% Coomassie blue solution (50% methanol, 50% H_2_O) with agitation at room temperature. A solution of 50% methanol, 40% H_2_O and 10% acetic acid was used to destain the gel.

### Preparation of human tau fractions

To generate the tau fractions used in ELISA and efficacy experiments, the protein was isolated from a human patient presenting with a mixture of AD/Pick’s disease pathology (National Disease Research Interchange, Philadelphia PA) as previously described [48, 49, 52]. The samples were selected based on their ability to induce toxicity in mixed cortical cultures. Isolation was carried out based on previously described methods. Briefly, the brain tissue was homogenized in a buffer containing 0.75 M NaCl, 1 mM EGTA, 0.5 mM MgSO_4_, and 100 mM 2-(*N*-morpholino) ethanesulfonic acid (pH 6.5). The homogenate was centrifuged at 11,000 x g for 20 minutes and this low-speed supernatant was incubated with 1% sarkosyl (from a 10% solution dissolved in PBS) for one hour at room temperature. Following the incubation period, the supernatant was centrifuged at 100,000 x g for 60 min. The supernatant containing the monomeric sarkosyl soluble tau was retained, and the pellet was washed with 1% sarkosyl. To generate the solubilized PHF fraction, the pellet was resuspended and then briefly heated to 37°C in 50 mM Tris-HCl buffer and dialyzed in PBS overnight. This PHF fraction contains soluble tau oligomers and the larger aggregated tau species.

### Primary culture

Mixed cortical cultures were prepared as described previously [52]. Briefly, culture plates were coated with poly-D-lysine for at least 3 h. Brains were collected from JNPL3 pups at postnatal day 0. The brainstem and meninges were removed, and the cortex/hippocampus was washed three times in modified Hank’s Balanced Salt Solution. The brains were then incubated with 0.5% trypsin for 20 min, followed by manual dissociation. Cultures were maintained in DMEM containing 20% serum, glucose and B27 supplement.

### PHF treatment

We tested the efficacy of each Ab in preventing PHF-induced toxicity and seeding as measured by GAPDH and total tau levels. Mixed cortical cultures were treated with 10 μg/ml of human-derived PHF and 10 μg/ml of one of the tau Abs using one of two dosing paradigms, which we have used to model extra- and intracellular mechanisms [48, 49, 52]. The first method mimics the extracellular blockage of tau spread. The tau and Abs were added to the culture at the same time (PHF + Ab) and allowed to incubate for 24 h. After this time, the media was exchanged, and the cells were collected after 5 days. In the second intracellular paradigm, we first incubated the cultures with PHF alone for 24 h, which allows the tau to be internalized by the neurons. Following several washes to remove any remaining extracellular PHF tau, we then incubated the cells in media containing the Ab for an additional 24 h, followed by washes to remove any remaining Abs, and collection for analysis 5 days later. As the PHF has already been taken up, in this method the Ab must also be internalized to be effective.

In all cultures, a set of day 0 control samples were collected prior to treatment. In each experiment a separate group of cells was also left untreated and collected with the experimental samples. The inclusion of the two control groups ensures that changes in GAPDH or tau are not simply the result of normal changes over time during culturing.

### Immunoblotting

Immunoblotting was conducted using methods previously described [47, 48, 51]. Cultures were washed with PBS and then lysed in 200 μL of RIPA buffer (50 mm Tris-HCl, 150 mm NaCl, 1 mm EDTA, 1% Nonidet P-40, 1X protease inhibitor cocktail (cOmplete^TM^, Roche), 1 mM NaF, 1 mM Na_3_VO_4_, 1 mM PMSF, 0.25% sodium deoxycholate, pH 7.4). A BCA assay was conducted, and protein concentration was normalized between samples. Aliquots of cell lysate were then added to the O+ loading buffer (62.5 mM Tris-HCl, 10% glycerol, 5% β-mercaptoethanol, 2.3% SDS, 1 mM EDTA, 1 mM EGTA, 1 mM NaF, 1 mM Na_3_VO_4_, 1 mM PMSF, 1X protease inhibitor cocktail (cOmplete^TM^, Roche), and boiled for 5 min. All samples prepared from the same animal and culture plate were run on the same gel, including both day 0 and day 5 untreated controls. Samples were run on a 12% polyacrylamide gel and then transferred to nitrocellulose membranes.

Blots were blocked with 5% milk in Tris buffered saline containing 0.1% Tween 20 (TBS-T) and then probed with Abs recognizing GAPDH (RRID: AB_10622025) and total tau (RRID: AB_10013724) at a 1:1000 dilution in Superblock^TM^ (ThermoFisher Scientific, Waltham, MA) overnight at 4°C. Membranes were washed three times in TBS-T and then incubated in HRP-conjugated rabbit secondary Ab (1:5000, RRID: AB_228341) for one hour at room temperature. Blots were then washed three times in TBS-T. Signal was visualized using a Fuji LAS-4000 and quantified with Image J (RRID:SCR_003070).

### CypHer5 labeling

For the uptake imaging experiments, all Abs were labeled with CypHer 5, a dye which fluoresces in acidic environments such as the endosomal/lysosomal system. The advantage of this marker is that it allows us to confirm that any fluorescence detected is in fact intracellular. Per manufacturer’s instructions (Cytiva) Abs were incubated with the dye in a solution of PBS and carbonate buffer for 1 h at room temperature. Abs were then dialyzed to remove unbound dye, and the Ab concentration was determined.

### Imaging

Cultures used for imaging were prepared as described above on glass bottom plates. Cells were incubated with 5 μg/ml of the labeled Abs for 1 h. Cultures were washed with PBS and fixed using 4% paraformaldehyde (PFA) and then imaged using an Axio Observer inverted confocal microscope at 20x magnification. Images were imported into ImageJ and the threshold was adjusted to isolate the fluorescence from the Abs. The percentage of pixels in each image with Ab signal was then determined.

### Enzyme-linked Immunosorbent Assay (ELISA)

We first determined the binding affinity for each of the Abs for four different tau peptides comprising different phosphorylation states of the 396/404 epitope:

non-phosphorylated peptide (RENAKAKTDHGAEIVYKSPVVSGDTSPRHL),
pSer396/pSer404 (TDHGAEIVYKpSPVVSGDTpSPRHL),
pSer396(TDHGAEIVYKpSPVVSGDTSPRHL),
or pSer404 (TDHGAEIVYKSPVVSGDTpSPRHL).

Plates were coated overnight at 4°C with one of the four peptides. Abs were serially diluted (33.0 - 0.0001 nM) and added to plate after 30 min of blocking at room temperature. The Abs were incubated for 2 h, after which the plates were washed and either HRP-conjugated anti-mouse or anti-human IgG was added at 1:5000. After 1 h of incubation with the secondary Abs, plates were washed again and developed using TMB peroxidase EIA reagent (ThermoFisher) and stopped by 2 N sulfuric acid. Absorbance at 450 nm was read on a BioTek Synergy 2 plate reader.

To test Ab binding to human tau, two types of ELISA were used. In the first type, plates were coated with tau from either the sarkosyl soluble fraction containing tau monomers, or the solubilized PHF fraction which contains oligomers and larger aggregates. One μg of each fraction was added per well and plates were incubated overnight at 4°C. Before the Abs were added, plates were washed and blocked for 30 min with Superblock^TM^. Abs were brought to the same concentration, and then serial dilutions were made. Abs were added to the plate and incubated for 2 h. Following this period, plates were washed again, and then an anti-mouse or anti-human secondary Ab (1:5000) was added for 1 h. Plates were washed and developed using TMB Peroxidase (ThermoFisher), and 2 M sulfuric acid to stop the reaction. Absorbance values were collected using a BioTek Synergy 2 plate reader.

For the second type, the competition ELISA, a single Ab dose was pre-incubated with increasing concentrations of solubilized PHF (0 – 200 ng) for 30 min before being added to the plate. After this period, the assay was carried out using the same methods as for the standard ELISA in plates coated with the solubilized PHF fraction.

### Fab fragment production and purification

The Fab fragment was prepared by papain digestion [36]. Briefly, IgG and papain (Worthington, Lakewood, NJ) were mixed at a 1:10 molar ratio in a buffer (20 mM Tris pH 6.8 and 100 mM NaCl) containing 20 mM cysteine hydrochloride (ThermoFisher) and 0.1 M EDTA pH 8.0. The reaction was incubated for 1 h at 37°C and was stopped by 10 mM iodoacetamide (Bio-Rad, Hercules, CA). The Fab fragments were then isolated from the Fc fragments using a HiTrap Protein A affinity column (Cytiva, DE). The Fab fragments were also cloned into VRC8400 vector by restriction enzymes EcoR1 and AfeI. The Fabs were expressed from 293 cells and purified by Ni-NTA column. Finally, the Fab fragments were further purified using size-exclusion chromatography. The monodispersed peak containing the soluble Fab fragments was collected and concentrated to more than 10 mg/mL for crystallization.

### Crystallization, data collection, and structure determination

Crystallization conditions were screened and optimized using the vapor diffusion hanging drop method. Well-diffracted crystals of Fab PHF-1/pSer396 complex (peptide sequence TDHGAEIVYK(pSer)PVVSGDTSPRHL) were obtained in a solution containing 16% polyethylene glycol 8000, 0.16 M magnesium acetate tetrahydrate, 0.08M sodium cacodylate trihydrate pH 6.5 and 20% glycerol. Well-diffracted crystals of Fab PHF-1 were obtained in a solution containing 26% polyethylene glycol 4000, 0.17 M ammonium sulfate and 15% glycerol. Well-diffracted crystals of Fab 9E were obtained in a solution containing 26% polyethylene glycol 8000, 0.085 M sodium cacodylate pH 6.5, 0.17 M ammonium sulfate and 15% glycerol. Well-diffracted crystals of Fab G10 were obtained in a solution containing 26% polyethylene glycol 8000, 0.17 M ammonium sulfate, and 15% glycerol. X-ray diffraction data were collected at 100 K at a wavelength of 0.979 Å at the NSLS II AMX beamline 17–1. The data sets were processed using the XDS software package and the structures determined by molecular replacement using initial models with high sequence similarity (**Table 4**) [59]. Multiple steps of refinement were carried out in COOT [60] and PHENIX [61]. Residues of the mAbs were numbered based on the Kabat scheme by using the web tool Abnum (www.bioinf.org.uk/abs/abnum/). The final structure analyses were carried out in ICM [62]. The antigen-Ab interactions were analyzed with the web tools PISA (www.ebi.ac.uk/msd-srv/prot_int/pistart.html). Figures were generated with Chimera [63] and PyMOL (pymol.org). The crystallization of the 4E6 Ab has been previously described [35].

### Statistics

Data was analyzed using GraphPad version 10.3.1 (RRID:SCR_002798). Peptide binding data was fit to a nonlinear regression using a one-site specific binding model. For uptake and efficacy experiments a one-way ANOVA was used and the difference between individual samples determined by Tukey’s multiple comparisons test. ELISA data was analyzed using a two-way ANOVA and Sidak’s multiple comparisons test. No data points were excluded from the analysis.

## Results

### Tau antibodies bind to the pSer396 and pSer404 epitope region, with 4E6 and 9E selective for pSer404 and G10 and PHF-1 selective for pSer396

To generate the 9E and G10 mAbs, ten mice were immunized using a DNA prime and protein boost immunization regimen (**Figure 1A, B**) [55]. The mice were first immunized with a DNA plasmid encoding the full-length tau protein, then the mice were split into two groups (n = 5) and boosted with the two different peptide immunogens (**Figure 1A**). Both groups had strong Ab responses (**Figure 1C**), and one animal (in Group 1) was selected for developing mAbs using the hybridoma method from its splenocytes by a commercial vendor. Two hybridoma cell lines, 9E and G10, were developed and the amino acid sequences of the mAbs were determined. Mouse mAbs were produced from hybridoma and the chimeric mAbs were recombinantly cloned into a mammalian expression vector, produced from 293F cells, and purified by protein G and protein A columns, respectively.

We then tested the binding of each murine and chimeric Ab to four peptides in their solid state, comprising the 396/404 epitope in different phosphorylation states (**Table 1**). We previously reported affinity data for the murine and chimeric 4E6 using solution phase methods, but this report is the first comparing solid state binding for all of the antibodies used herein [35, 49]. Murine 4E6 was the only one of the murine Abs to bind to the non-phosphorylated peptide with a K_D_ of 0.058 nM (**Figure 2A**). All the murine Abs bound to the pSer396/pSer404 peptide (K_D_ = 0.00467 (4E6), 0.0120 (9E), 0.0207 (G10) and 0.0157 nM (PHF-1), **Figure 2B**). Murine G10 and PHF-1 bound to the pSer396 peptide (K_D_ = 0.0338 and 0.0153 nM, **Figure 2C**) while 4E6 and 9E bound only at the highest concentrations and did not reach a plateau. Conversely, murine 4E6 and 9E bound to the pSer404 peptide (K_D_ = 0.00556 and 0.00850 nM, **Figure 2D**) while G10 and PHF-1 bound poorly to it.

**Figure 2.**
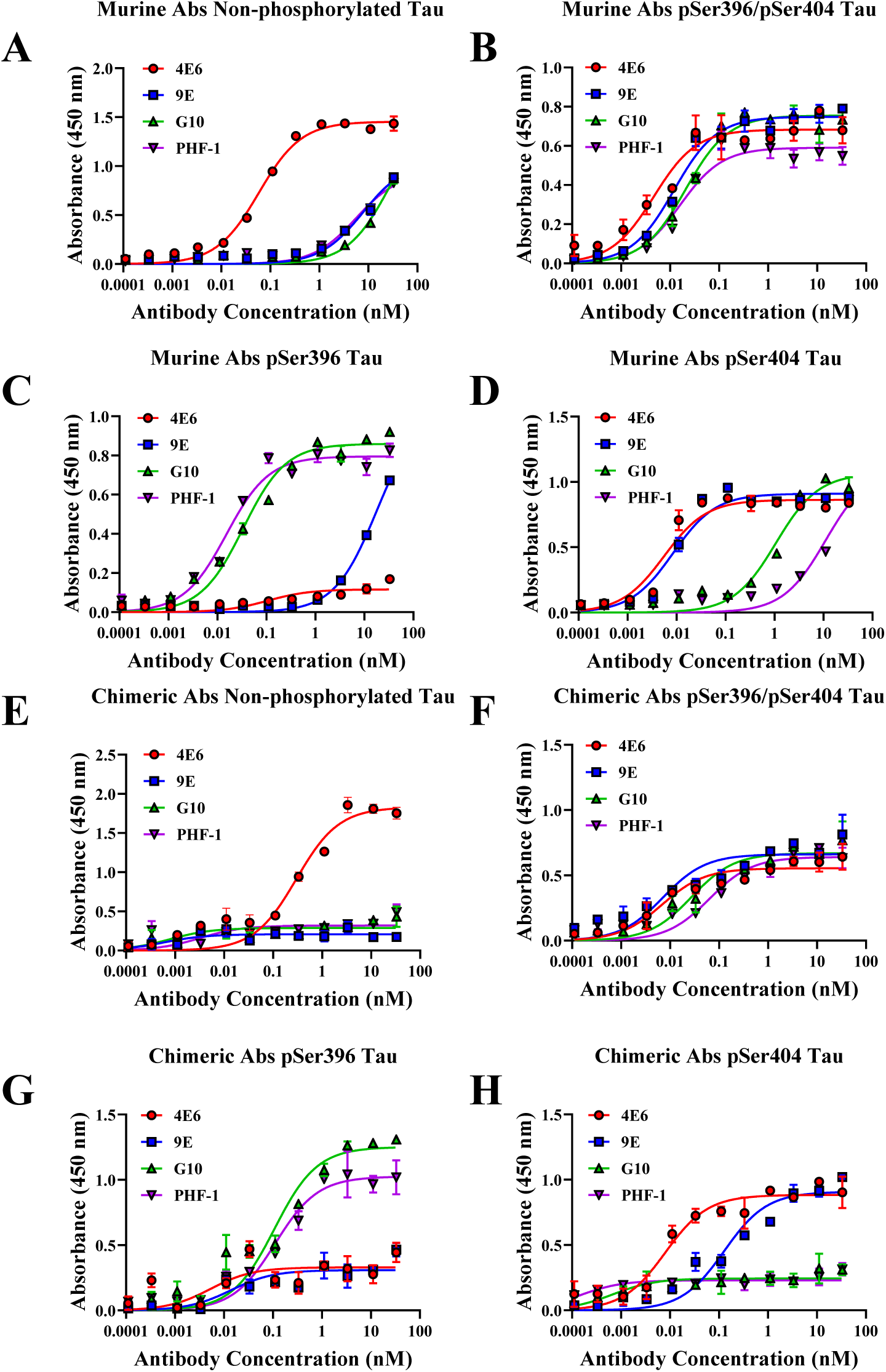
Murine and chimeric antibodies (Abs) show differential binding to tau peptides comprising the 396/404 epitope. Abs were added to plates coated with four different peptides comprising the different phosphorylation states of the 396/404 epitope of tau at concentrations from 33.0 - 0.0001 nM. **A-D.** Murine antibody binding to plates coated with peptides containing non-phosphorylated 396/404, pSer396/pSer404, pSer396, and pSer404 tau, respectively. **E-H**. Chimeric antibody binding to plates coated with peptides containing non-phosphorylated 396/404, pSer396/pSer404, pSer396, and pSer404 tau, respectively.

**Table 1.** K_D_ values for murine and chimeric antibody binding to pSer396/pSer404 tau peptides.

| **$K_D$ (nM)** | **non-p tau** | **p396/404 tau** | **p396 tau** | **p404 tau** |
| --- | --- | --- | --- | --- |
| **Murine 4E6** | 0.058 | 0.00467 | NB | 0.00556 |
| **Murine 9E** | NB | 0.0120 | NB | 0.00850 |
| **Murine G10** | NB | 0.0207 | 0.0338 | NB |
| **Murine PHF-1** | NB | 0.0157 | 0.0153 | NB |
| **Chimeric 4E6** | 0.299 | 0.00691 | NB | 0.008 |
| **Chimeric 9E** | NB | 0.00716 | NB | 0.128 |
| **Chimeric G10** | NB | 0.0275 | 0.108 | NB |
| **Chimeric PHF-1** | NB | 0.0635 | 0.0111 | NB |

non-phosphorylated peptide: RENAKAKTDHGAEIVYKSPVVSGDTSPRHL
pSer396/pSer404: TDHGAEIVYKpSPVVSGDTpSPRHL
pSer396: TDHGAEIVYKpSPVVSGDTSPRHL
pSer404: TDHGAEIVYKSPVVSGDTpSPRHL
Values were determined using ELISA data fit to a one-site specific binding nonlinear curve.

The binding pattern was the same for the chimeric Abs with 4E6 being the only one to bind to the non-phosphorylated peptide (K_D_ = 0.2989 nM, **Figure 2E**). As with the murine variants, all four chimeras bound to the pSer396/pSer404 peptide (K_D_ = 0.00691 (4E6), 0.00716 (9E), 0.0275 (G10) and 0.0635 nM (PHF-1), **Figure 2F**). Chimeric G10 and PHF-1 bound to the pSer396 peptide (K_D_ = 0.108 and 0.111 nM), while 4E6 and 9E bound poorly to it (**Figure 2G**). 4E6 and 9E chimeras bound to pSer404 (K_D_ = 0.008 and 0.128 nM), whereas G10 and PHF-1 bound poorly to it (**Figure 2H**). In conclusion, we found that 9E was pSer404 dependent while G10 was pSer396 dependent.

### Chimerization changes antibody binding to human tau

In our previous publications, we have found that changing Ab subclass or undergoing chimerization can impact Ab binding to human tau [48, 52]. This binding has also been shown to relate to efficacy more closely than binding to the peptide antigen [48, 49, 52]. To measure binding, each murine/chimera pair were assayed for binding to sarkosyl soluble monomeric tau and solubilized PHF using standard solid phase ELISA assay, and preference for soluble tau in a competitive ELISA.

In the first set of ELISA experiments, plates were coated with 1 μg per well of the sarkosyl soluble tau fraction. As in previous experiments, we observed clear difference in binding between murine and chimeric 4E6 (Ab, concentration, and interaction effects, two-way ANOVA, p < 0.0001 for all, **Figure 3A**). The chimeric 4E6 bound stronger to this tau fraction than its murine version from 1/100 – 1/312.5K dilutions (p = 0.001- < 0.0001). The murine and chimeric 9E differed also in concentration and interaction effects (p < 0.0001 for both, **Figure 3B**), but contrastingly chimeric 9E bound less to this tau fraction compared to the murine variant at the 1/100 and 1/500 dilutions (p < 0.0001, = 0.002). The G10 versions differed in Ab and concentration effects in their binding to this fraction (p < 0.0001, = 0.0004, **Figure 3C**). Chimeric G10 binding to sarkosyl soluble tau was higher from 1/100 – 1/62.5K dilutions (p = 0.04 – 0.0009, **Figure 3C**). Likewise, the PHF-1 versions differed in Ab and concentration effects (p < = 0.0001, = 0.002, **Figure 3D**), with the chimeric PHF-1 binding better to this tau fraction at dilutions from 1/100 – 1/12.5K (p = 0.04 – 0.003).

**Figure 3.**
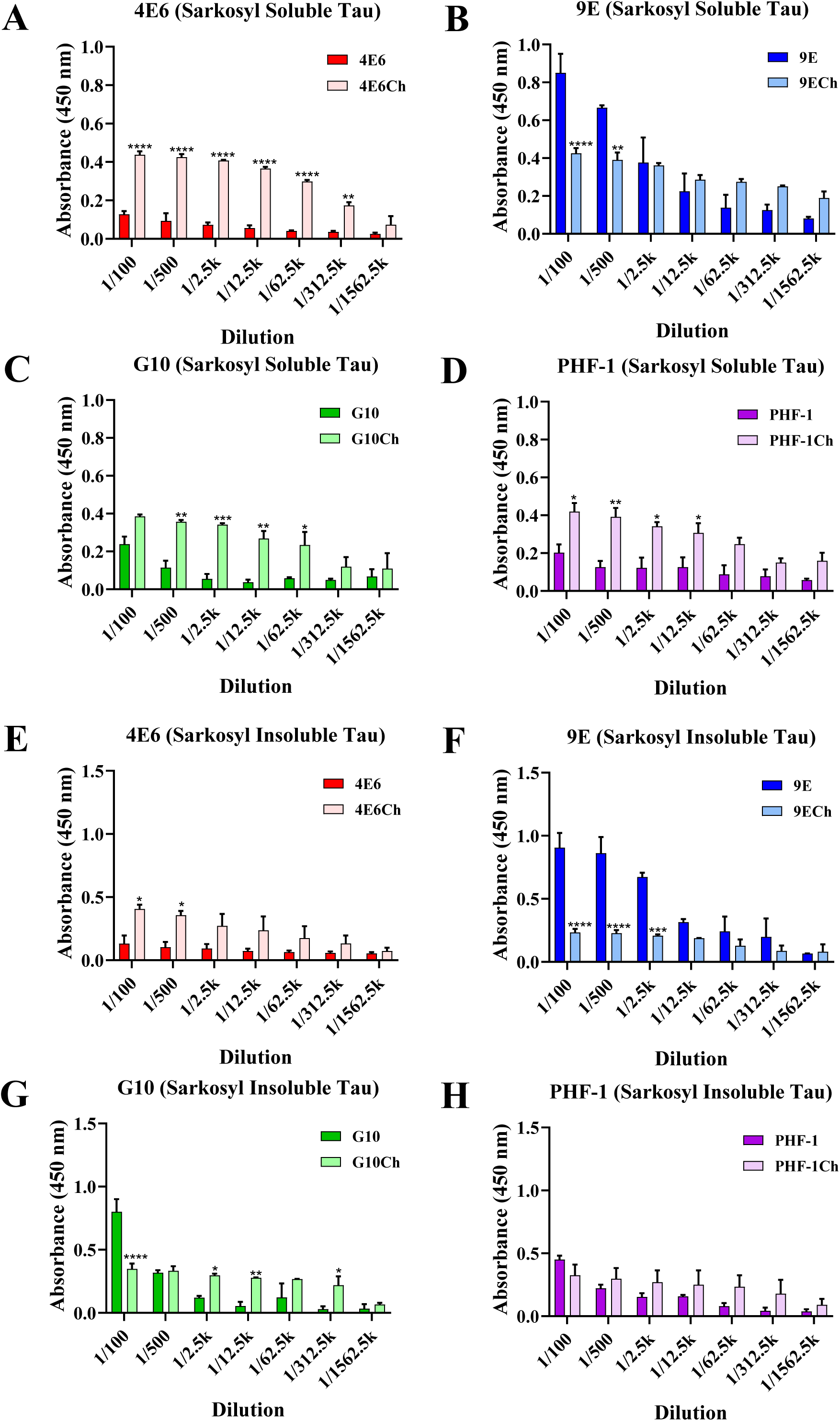
Chimerization changes antibody binding to human tau. Plates were coated with 1 μg per well of either sarkosyl soluble tau (**A-D**), or solubilized PHF tau derived from the sarkosyl insoluble fraction (**E-H**). Plates were blocked and then incubated with serial dilutions of murine and chimeric 4E6, 9E, G10, or PHF-1 for two hours. **A**. In plates coated with sarkosyl soluble tau, chimeric 4E6 had higher binding from 1/100 – 1/312.5K dilutions (p = 0.001 - < 0.0001). **B**. Chimeric 9E had lower binding compared to the murine variant at the 1/100 and 1/500 dilutions (p < 0.0001, = 0.002, respectively). **C**. Chimeric G10 binding to sarkosyl soluble tau was higher from 1/100 – 1/62.5K dilutions (p = 0.04 – 0.0009). **D**. Chimeric PHF-1 binding was higher at dilutions from 1/100 – 1/12.5K (p = 0.04 – 0.003). **E**. When the plate was coated with tau from the sarkosyl insoluble fraction, chimeric 4E6 had higher binding at the 1/100 and 1/500 dilutions (p = 0.05 – 0.03). **F**. Chimeric 9E had lower binding at the 1/100-1/2.5K dilutions (p = 0.0001 - < 0.0001). **G**. Chimeric G10 had a lower signal at 1/100 and higher at 1/2.5K and 1/12.5K (p < 0.0001, = 0.02, and = 0.003 respectively). **H**. There were no significant differences in absorbance between murine and chimeric PHF-1. * p ≤ 0.05, ** p ≤ 0.01, *** p ≤ 0.001, **** p < 0.0001

The same approach with solubilized PHF-enriched tau fraction revealed similar differences in binding between the murine and chimeric version for 4E6, 9E and G10 but not for PHF-1. Specifically, for 4E6, its two versions differed in their binding (Ab and concentration effects p = 0.002, = 0.04, **Figure 3E**), with higher binding at the 1/100 and 1/500 dilutions seen with chimeric 4E6 (p = 0.05 – 0.03). The 9E versions differed as well in their binding to this tau fraction (Ab, concentration, and interaction effects, p < 0.0001 for all, **Figure 3F**). Compared to its murine variant, chimeric 9E had lower binding at the 1/100-1/2.5K dilutions (p = 0.0001 - < 0.0001). The same factors differed as well for the G10 versions (Ab, concentration, and interaction effects, p = 0.02, < 0.0001, < 0.0001, **Figure 3G**). The chimeric G10 bound less at the 1/100 dilution, but higher at 1/2.5K and 1/12.5K (p < 0.0001, = 0.02, and = 0.003). In contrast, the two PHF-1 versions did not differ in their binding to the PHF tau fraction (**Figure 3H**).

We then conducted a competition ELISA using a single concentration of each Ab. Plates were coated with 1 μg/well with tau from the solubilized PHF fraction. Aliquots were preincubated with increasing doses of solubilized PHF for 30 minutes before being added to the plate. In previous experiments we have found that binding to tau in solution, and thus showing reduced binding to the tau coated on the assay plate, to be an indicator of Ab efficacy in culture [48, 49, 52]. The two 4E6 versions differed in their binding (Ab, PHF dose, and interaction effects (p < 0.0001 for all, **Figure 4A**). The murine version had a dose-dependent decrease in binding, whereas chimeric 4E6 did not (23 – 71% reduction, p < 0.0001 for all). The 9E versions differed as well (Ab and PHF dose effects (p = 0.005, < 0.0001, **Figure 4B**). The murine variant showed a dose-dependent decreased binding at the three highest PHF doses (40 – 69% reduction, p = 0.0003 - < 0.0001) and chimeric 9E at the highest two doses (44 – 54% reduction, p = 0.0001, < 0.0001). The G10 versions differed as well (Ab and PHF dose effects, p < 0.0001 for both, **Figure 4C**). The murine G10 bound less to the solid phase tau at the highest PHF dose in solution (49 % reduction, p = 0.005), and the chimeric G10 dose dependently at the three highest PHF doses (47 – 64% reduction, p = 0.008 – 0.0003). Lastly, the PHF versions differed as well in the binding profile (Ab, PHF dose, and interaction effects, p < 0.0001, = 0.0002, = 0.002, **Figure 4D**). Only murine PHF-1 had a dose-dependent reduced binding to the plate at the four highest PHF doses (27 – 51% reduction, p = 0.04 - < 0.0001).

**Figure 4.**
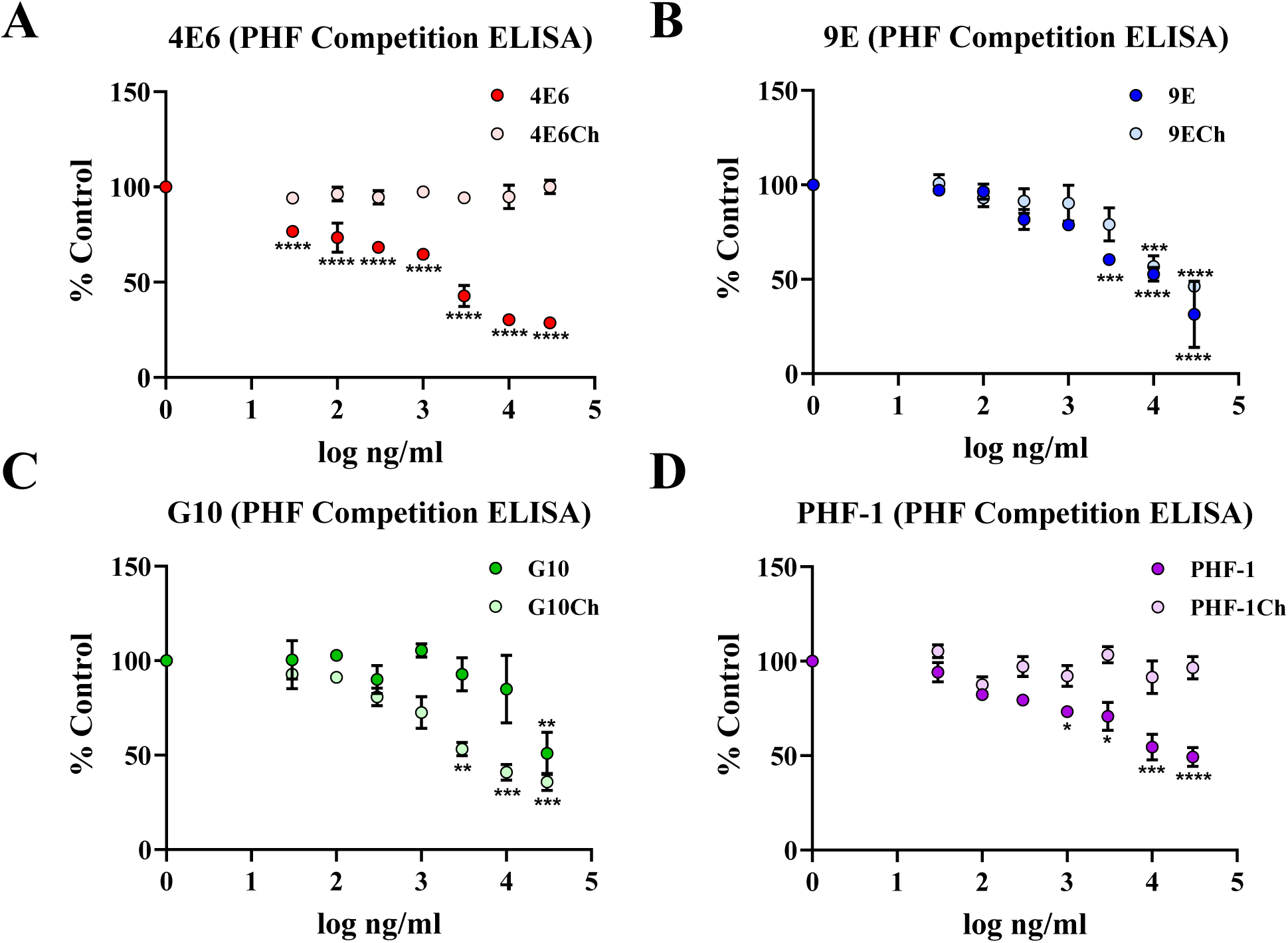
Human chimerization alters selectivity for soluble tau species. A single concentration of each antibody was preincubated with increasing doses of solubilized PHF before being added to the plate. **A**. Murine 4E6 showed reduced binding to the plate at every PHF concentration used (p < 0.0001) while chimeric 4E6 did not. **B**. Murine 9E showed decreased binding at the three highest PHF concentrations (40 – 69% reduction, p = 0.0003 - < 0.0001) and chimeric 9E at the highest two (44 – 54% reduction, p = 0.0001, < 0.0001). **C.** Murine G10 had reduced binding at the highest PHF concentration (49 % reduction, p = 0.005), and chimeric G10 at the three highest (47 – 64% reduction, p = 0.008 – 0.0003). **D**. Murine PHF-1 had reduced binding to the plate at the four highest tau concentrations (27 – 51% reduction, p = 0.04 - < 0.0001). * p ≤ 0.05, ** p ≤ 0.01, *** p ≤ 0.001, **** p < 0.0001

We also conducted a competition ELISA for the 6B2 and 8B2 Abs. The assay was conducted using the same methods as above. No reductions in binding to the plate were observed with 6B2 at any PHF doses (**Figure S1**). 8B2 however, had dose-dependent lower binding to the plate at all but the lowest PHF dose (37-50% reduction, p = 0.001 - < 0.0001, **Figure S1**).

### Chimerization increases isoelectric point (IEP)

Aliquots from each of the murine/chimera pairs were run on an isoelectric focusing gel and detected with Coomassie stain. In all cases, the change from murine to human constant domains resulted in an increase to a higher IEP for Ab charge (**Figure 5**, **Table 2**). Specifically, 4E6 went from 6.1 – 6.5 to 8.0., 9E from 7.8 to > 9.6, G10 from 7.1 – 7.25 to > 9.6, and PHF-1 from 7.5 – 7.8 to > 9.6. In our previous study [48], murine and chimeric 4E6 had IEPs of 6.5 and 9.6 respectively. Although the values from this experiment are lower, they are still consistent with the overall pattern of increased IEP following chimerization. Together with the binding data, these results further highlight that the change from mouse to human forms can alter Ab properties [48].

**Figure 5.**
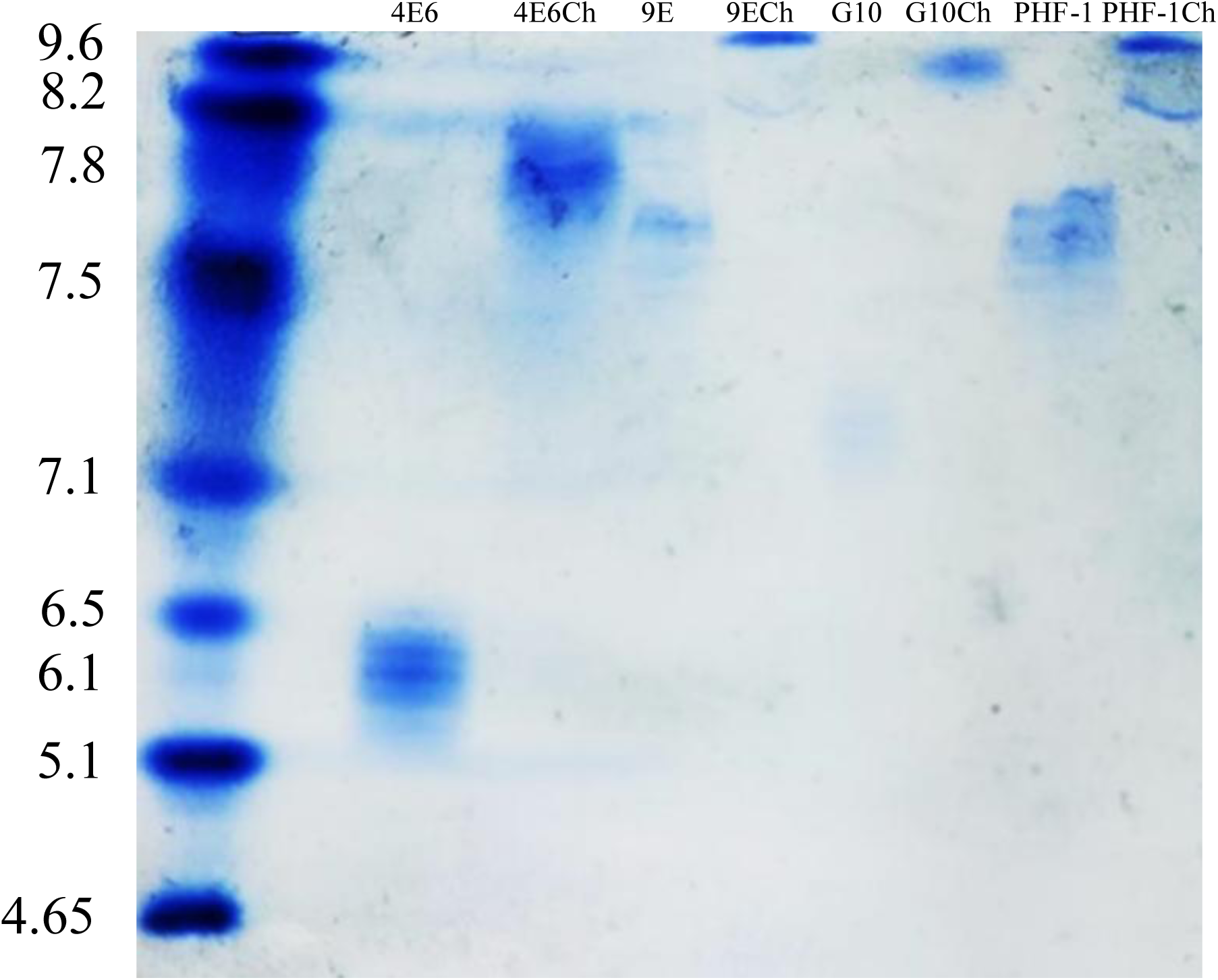
Chimerization increases isoelectric point (IEP). For all of the Abs tested, chimerization resulted in an increase in the isoelectric point. 4E6 from 6.1-6.5 to 8.0, for 9E 7.8 to > 9.6, for G10 7.1-7.25 to 9.6, for PHF-1 7.5-7.8 to > 9.6.

**Table 2.** Isoelectric points of murine and chimeric antibodies.

|  | **Murine** | **Chimera** |
| --- | --- | --- |
| 4E6 | 6.1 - 6.5 | 8.0 |
| 9E | 7.8 | > 9.6 |
| G10 | 7.1 - 7.25 | = 9.6 |
| PHF-1 | 7.5 - 7.8 | > 9.6 |

Values were obtained by running antibodies on a precast IEF gel (BioRad). Chimerization increased IEP for all antibodies tested.

### Antibody (Ab) efficacy

The efficacy of each murine/chimera pair was tested in mixed cortical cultures containing neurons and glia using two different dosing methods.

In the first paradigm, 10 μg/mL PHF and 10 μg/mL of Ab were added to the culture media at the same time (PHF + Ab). This method simulates extracellular interactions between tau and the Abs, such as blocking uptake into neurons and promoting microglial phagocytosis and does not require the Ab to be internalized by neurons.

In the second paradigm, mixed cortical cultures were incubated with 10 μg/mL PHF for 24 hours. The media was then removed, the cells washed thoroughly to remove any PHF that had not been taken up into cells, and media added with 10 μg/mL of one of the murine or chimeric Abs. Under these conditions, the PHF has already been internalized once the Ab is added, and to bind and neutralize it the Ab must be internalized as well. In previous experiments, we have shown that blocking or reducing Ab uptake impairs efficacy in this dosing paradigm [48, 49, 52].

For both conditions, a set of untreated control cells were collected on the day of treatment to control for any changes that may naturally occur in the cultures. The treated cells and a second set of controls were collected 96 hours after tau/Ab exposure.

### Extracellular efficacy: In the PHF + Ab dosing paradigm, murine 4E6, 9E, PHF-1, and chimeric G10 prevent toxicity, while all Abs except chimeric 9E and chimeric PHF-1 prevent tau seeding

Toxicity was assessed using immunoblotting for GAPDH (**Figure 6A**). There was an overall treatment effect (p < 0.0001, **Figure 6B**). PHF alone was toxic, with GAPDH levels at 30% of controls (p < 0.0001). This PHF-induced toxicity was prevented by murine 4E6, murine 9E, chimeric G10, and murine PHF-1 with GAPDH levels at 120, 85, 160, and 71% control values (p < 0.0001, < 0.0001, < 0.0001, 0.002 respectively). In contrast, chimeric 4E6, 9E, murine G10 and chimeric PHF-1 did not prevent PHF toxicity with GAPDH levels at 42, 51, 35.0 % control values (p < 0.0001, for all relative to untreated cells) **Figure 6B**).

**Figure 6.**
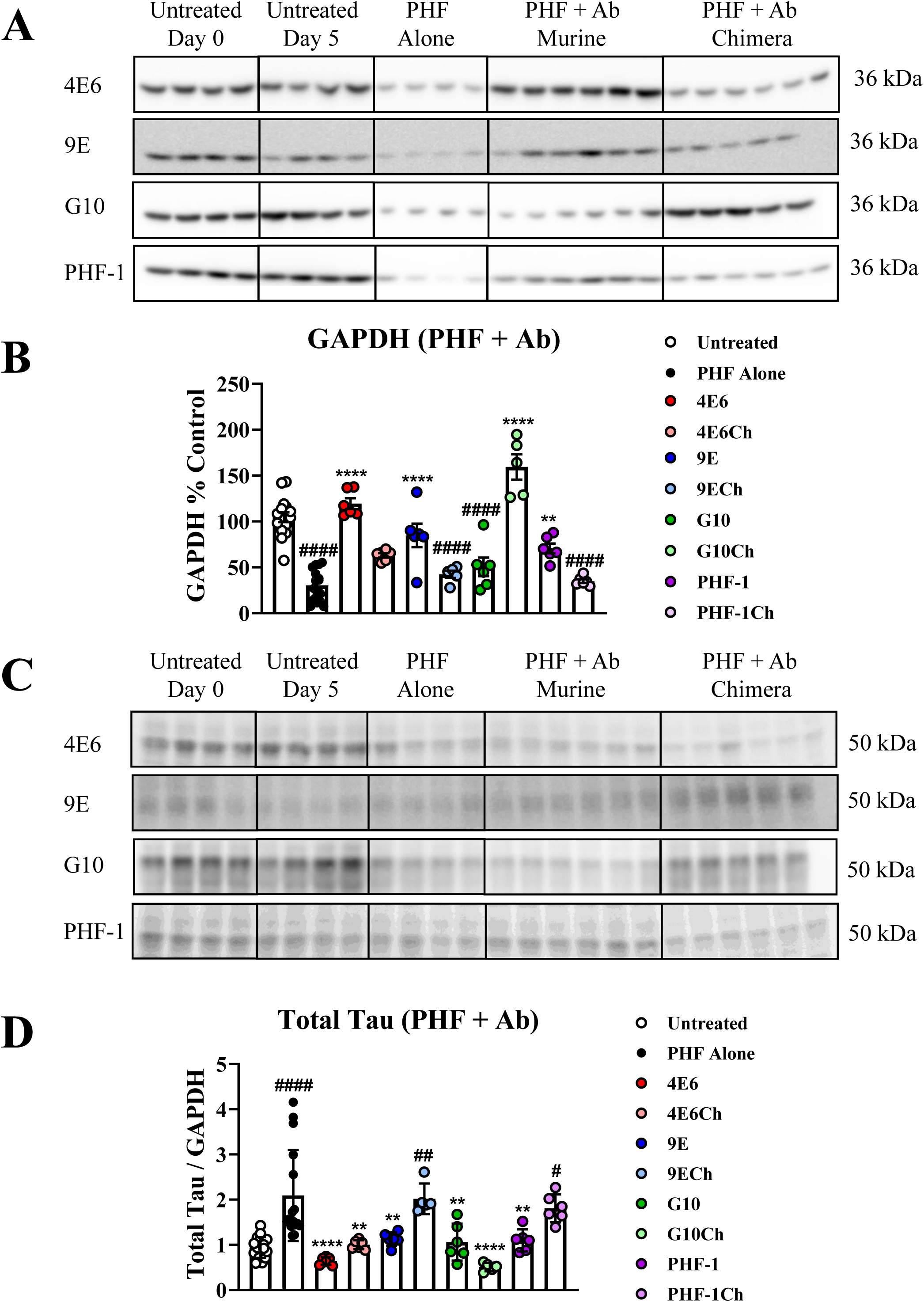
Extracellular efficacy: In the PHF + Ab dosing paradigm, murine 4E6, 9E, PHF-1, and chimeric G10 prevent toxicity, while all Abs except chimeric 9E and chimeric PHF-1 prevent tau seeding. Mixed cortical cultures were incubated with 10 μg/mL PHF and 10 μg/mL of antibody added to the culture at the same time. In this dosing method the antibody binds to tau in the extracellular space and can prevent neuronal uptake or facilitate phagocytosis by microglia. Cells were collected 96 hours after tau and antibody treatment. **A**. Immunoblots showing GAPDH levels in the treated cells. **B**. A one-way ANOVA showed a treatment effect on GAPDH levels in the cultures (p < 0.0001). Exposure to PHF alone resulted in toxicity (GAPDH 30 % of control values, p < 0.0001). Chimeric 9E, murine G10, and chimeric PHF-1 also had lower GAPDH relative to untreated controls (42, 51, and 35% of controls (p < 0.0001 for all). Murine 4E6, murine 9E, chimeric G10, and murine PHF-1 prevented PHF-induced toxicity and had higher GAPDH levels than samples exposed to PHF alone (119, 85, 159, and 70% of controls (p < 0.0001, < 0.0001, < 0.0001, and = 0.002 respectively). **C**. Immunoblots showing total tau levels in the treated cultures. **D**. An overall treatment effect was seen by one-way ANOVA (p < 0.0001). PHF alone increased total tau compared to control (Tau/GAPDH ratio 2.1, p < 0.0001). Chimeric 9E and PHF-1 also had higher tau levels than untreated controls (Tau/GAPH ratio 2.0 and 1.8, p = 0.005, = 0.03). Lower tau compared to PHF alone was seen in cultures cotreated with murine and chimeric 4E6 (Tau/GAPDH ratio 0.64 and 1.0, p < 0.0001, = 0.002), murine 9E (Tau /GAPDH ratio 1.1, p = 0.008), both forms of G10 (Tau/GAPDH ratio 1.1 and 0.51, p = 0.004, < 0.0001), and murine PHF-1 (Tau/GAPDH ratio 1.1, p = 0.006) # p ≤ 0.05, ## p ≤ 0.01, ### p ≤ 0.001, #### p < 0.0001 **** p < 0.0001

Samples were then assayed for total tau levels and normalized for GAPDH to account for any cell death or toxicity following PHF exposure (**Figure 6C**). As with the GAPDH levels, there was an overall treatment effect (p < 0.0001, **Figure 6D**). Incubation with PHF alone increased total tau compared to controls (Tau/GAPDH ratio 2.1, p < 0.0001). Both murine and chimeric 4E6 prevented the PHF-induced increase in tau (Tau/GAPDH ratio 0.64, 1.0, p < 0.0001, 0.002), as did murine 9E (Tau /GAPDH ratio 1.1, p = 0.008), murine and chimeric G10 (Tau/GAPDH ratio 1.1 and 0.51, p = 0.004, < 0.0001), and murine PHF-1 (Tau/GAPDH ratio 1.1, p = 0.006). In contrast, chimeric 9E or chimeric PHF-1 did not prevent the PHF-induced increase in tau (Tau/GAPDH ratio 2.0, 1.8, p = 0.005, 0.03 compared to controls).

In addition, we also compared the efficacy of two other Abs against the Ser396/Ser404 region, 6B2 and 8B2. In previous experiments, 6B2 has shown utility as an imaging agent but was not effective against tau either in culture or in vivo [46, 48, 64], whereas 8B2 has shown efficacy under such conditions [50]. Mixed cultures were treated using the same PHF + Ab dosing paradigm and collected 96 hours following treatment. As above, cells were assayed for levels of GAPDH and total tau (**Figure S2A**). There was an overall treatment effect (p < 0.0001, **Figure S2B**). As above, PHF alone was toxic (41% of controls, p < 0.0001) and its toxicity was prevented by 8B2 (141% of controls, p < 0.0001) but not by 6B2 (57% of controls, p < 0.0003).

As with the toxicity data, there was an overall treatment effect on total tau levels (p < 0.0001, **Figure S2C**). PHF alone increased total tau (Tau/GAPDH ratio 2.0, p = 0.0002), and 8B2 prevented this increase (Tau/GAPDH ratio 0.59, p < 0.0001 compared to PHF alone), whereas 6B2 was ineffective (Tau/GAPDH ratio 1.5, p = 0.009).

### Intracellular efficacy: In the PHF → Ab dosing paradigm, murine 4E6, both 9E variants, and chimeric G10 prevent tau toxicity and seeding

As above, toxicity was measured by GAPDH levels (**Figure 7A**). There was an overall treatment effect under these conditions (p < 0.0001, **Figure 7B**). Again, PHF alone induced toxicity resulting in decreased GAPDH levels relative to untreated samples (46 % control values, p < 0.0001). This toxicity was prevented with murine 4E6, both forms of 9E and chimeric G10 (101, 95, 99, 188 % untreated control values, p = 0.0003, 0.002, 0.0004, < 0.0001 respectively compared to PHF alone), whereas chimeric 4E6, murine G10, and both PHF-1 variants were ineffective (53, 58, 51, 25, p = 0.002, = 0.007, 0.0007, < 0.0001, respectively, compared to untreated controls).

**Figure 7.**
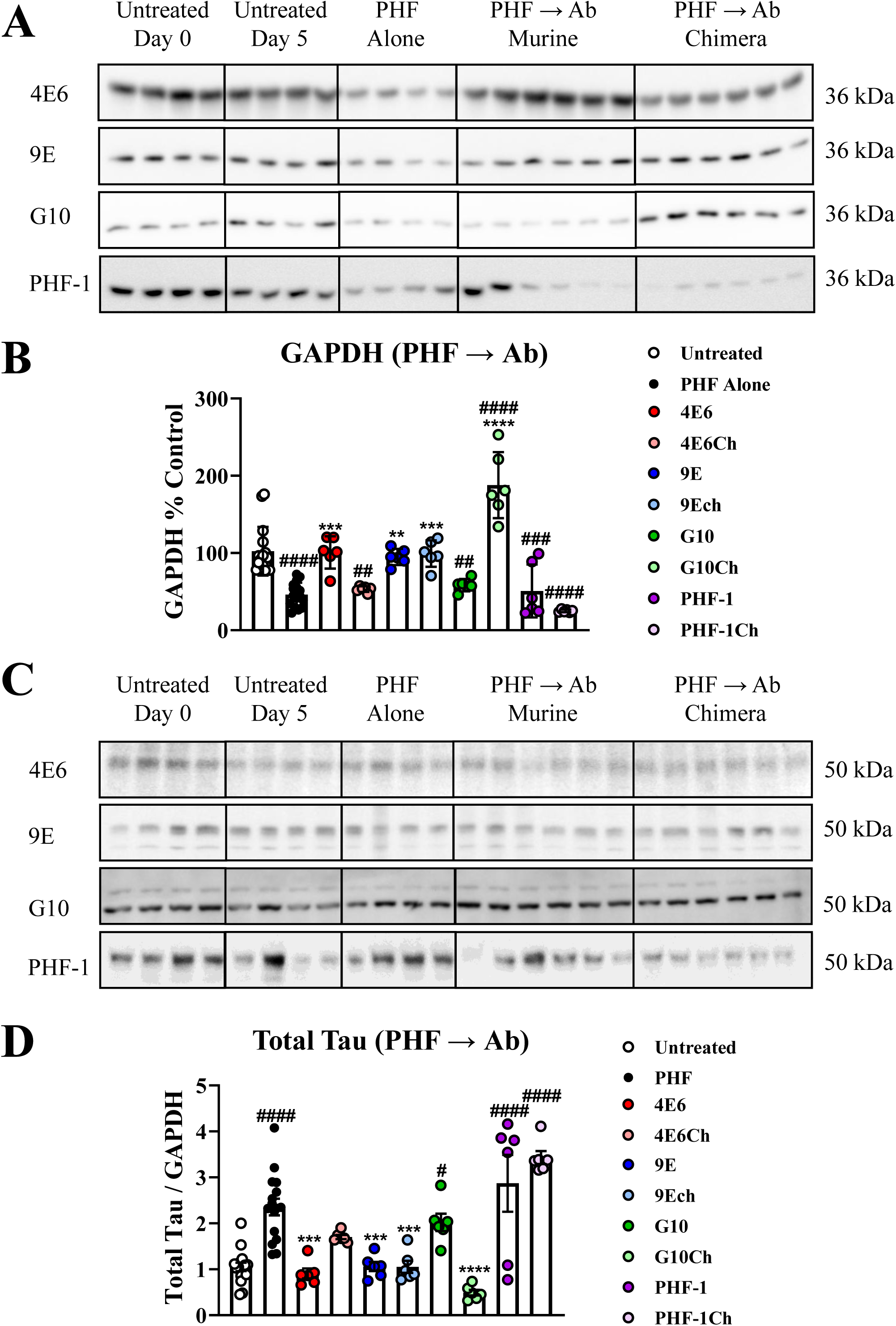
Intracellular efficacy: In the PHF → Ab dosing paradigm, murine 4E6, both 9E variants, and chimeric G10 prevent tau toxicity and seeding. Mixed cortical cultures were incubated with 10 μg/mL PHF, followed 24 hours later by 10 μg/mL of antibody. In this dosing paradigm the PHF is internalized, and the antibody must be taken up as well to be effective. Cells were collected 96 hours after antibody treatment. **A**. Immunoblots showing GAPDH levels in the treated cells. **B**. A one-way ANOVA showed a treatment effect on GAPDH levels in the cultures (p < 0.0001). Exposure to PHF alone resulted in toxicity relative to untreated control cells (46 % of control values, p < 0.0001). Chimeric 4E6, murine G10, and both murine and chimeric PHF-1 also had lower GAPDH relative to untreated controls (53, 58, 51, and 25 % of control values, p = 0.002, = 0.007, = 0.0007, and < 0.0001). Murine 4E6, murine and chimeric 9E, and chimeric G10 prevented the PHF-induced toxicity and led to GAPDH levels higher than samples exposed to PHF alone (101, 95, 99, and 188% of control values, p = 0.0003, = 0.002, 0.0004, < 0.0001 respectively). **C**. Immunoblots showing total tau levels in the treated cultures. **D**. An overall treatment effect was seen by one-way ANOVA (p < 0.0001). PHF alone increased total tau compared to control (Tau/GAPDH ratio 2.4, p < 0.0001). Murine G10 and both forms of PHF-1 were ineffective in decreasing these higher tau levels (Tau/GAPDH ratio 2.0, 2.9, and 3.4 p = 0.03, < 0.0001, < 0.0001). Lower tau compared to PHF alone was seen in cultures cotreated with murine 4E6, murine and chimeric 9E, chimeric G10 (Tau/GAPDH ratio 0.9, 1.1, 1,1, 0.5, p = 0.0001, 0.0008, 0.0008, < 0.0001). # p ≤ 0.05, ## p ≤ 0.01, ### p ≤ 0.001, #### p < 0.0001 * p ≤ 0.05, ** p ≤ 0.01, *** p ≤ 0.001, **** p < 0.0001

Samples were also probed for total tau (**Figure 7C**), and the values normalized for GAPDH levels. There was an overall treatment effect (p < 0.0001, **Figure 7D**). PHF alone increased tau levels relative to untreated controls (Tau/GAPDH ratio 2.4, p < 0.0001). This increase was prevented with murine 4E6, both forms of 9E, and chimeric G10 (Tau/GAPDH ratio 0.9, 1.1, 1.1, 0.5, p = 0.0001, 0.0008, 0.0008, < 0.0001), whereas murine G10, and both murine and chimeric PHF-1 were ineffective (Tau/GAPDH ratio 2.0, 2.9, 3.4, p = 0.03, < 0.0001, < 0.0001, compared to untreated controls). Intracellular tau was not significantly increased in cells treated with chimeric 4E6, but tau was also not lower when compared to PHF alone.

We also tested 6B2 and 8B2 by the PHF → Ab paradigm (**Figure S2D)**, with an overall treatment effect for toxicity (p = 0.004, **Figure S2E**). As before, PHF alone lowered GAPDH levels (69.6% of controls, p = 0.04) and its toxicity was prevented by 8B2 (141.2% control values, p < 0.0001), whereas 6B2 was ineffective (53.0% of controls, p = 0.003). PHF also increased tau levels (Tau/GAPDH ratio 2.4, p = 0.003, compared to untreated controls, **Figure S2F**), which was prevented by 8B2 (Tau/GAPDH ratio 0.59, p = 0.02, compared to PHF alone), whereas 6B2 was ineffective (Tau/GAPDH ratio 2.2, p = 0.05, compared to untreated controls).

Together these data indicate that chimerization can alter Ab efficacy, in some cases rendering it ineffective, and in others enhancing its efficacy (See **Table 3** for a summary of Ab efficacy). Prior results by us and others with modified Abs have shown that changes to the constant Ab regions can impact antigen binding and efficacy, although the variable regions remain the same [48, 52, 64–76]. Thus, when changing subclass or species, Abs should be retested to ensure that their efficacy remains intact. Further, as evident by chimeric 4E6 and murine G10, efficacy should not be measured solely by prevention of tau seeding, as these versions of the Abs prevented tau seeding in the PHF + Ab paradigm but not PHF-induced toxicity.

**Table 3.** Efficacy of murine and chimeric tau antibodies.

|  | **PHF + Ab**<br>**Toxicity** | **PHF + Ab**<br>**Seeding** | **PHF → Ab**<br>**Toxicity** | **PHF → Ab**<br>**Seeding** |
| --- | --- | --- | --- | --- |
| **4E6** |  |  |  |  |
| **Murine** | **** | **** | *** | *** |
| **Chimera** | - | ** | - | - |
| **9E** |  |  |  |  |
| **Murine** | **** | ** | ** | *** |
| **Chimera** | - | - | *** | *** |
| **G10** |  |  |  |  |
| **Murine** | - | ** | - | - |
| **Chimera** | **** | **** | **** | **** |
| **PHF-1** |  |  |  |  |
| **Murine** | ** | ** | - | - |
| **Chimera** | - | - | - | - |

Murine and chimeric antibodies differ in their ability to prevent PHF-induced toxicity and seeding in extracellular (PHF + Ab) and intracellular (PHF → Ab) dosing paradigms.
\* indicates significance relative to PHF alone samples.

### Murine and chimeric antibodies show similar levels of cellular uptake

In previous studies, we have found that Ab efficacy is linked to neuronal uptake in the PHF → Ab paradigm, and further that chimerization can drastically change an Ab’s ability to enter cells [48]. To examine this further here, each Ab was labeled with CypHer5 a dye that fluoresces in acidic compartments such as the endosomal/lysosomal system. Mixed cortical cultures were incubated with 5 μg/ml of the labeled Abs. In these experiments, we measured the uptake into all cell types, but in previous experiments we have found that the majority of uptake is neuronal [44]. Cultures were then fixed using 4% paraformaldehyde (PFA) and images were collected using an AxioObserver confocal microscope (**Figure 8A-H**). Images were imported into Image J and the percentage of pixels in each containing fluorescent signal was determined (**Figure 8I**). All Abs were internalized to a similar degree with no significant differences between the groups. In previously published experiments we found reduced uptake of chimeric 4E6 relative to its murine form [48], while in these experiments we saw no such difference. At the time of the previous publication, chimeric 4E6 was generated using both 293F and 293S cells [35, 48], and these two cell lines produce Abs with different glycosylation profiles. The glycosylation profile of an aliquot of the chimeric 4E6 used in the published efficacy experiments was examined and we confirmed that it was generated in 293F cells. However, based on the differences in uptake seen in the current results, we hypothesized that CypHer5 labeling may have been performed using the product of the 293S line. This sample was unavailable for testing, thus we generated new 4E6 chimeras from 293S and 293F cells (**Figure S3**) and labeled them with CypHer5. Cultures were incubated for 1 hour with either 1 or 2 μg/ml of the Ab (**Figure S4A**) and then fixed with 4% PFA and imaged as described above. The percentage of pixels per image containing the fluorescent signal was determined for each. At both doses, uptake of the 293S Abs was lower than the 293F version (p < 0.0001, **Figure S4B**).

**Figure 8.**
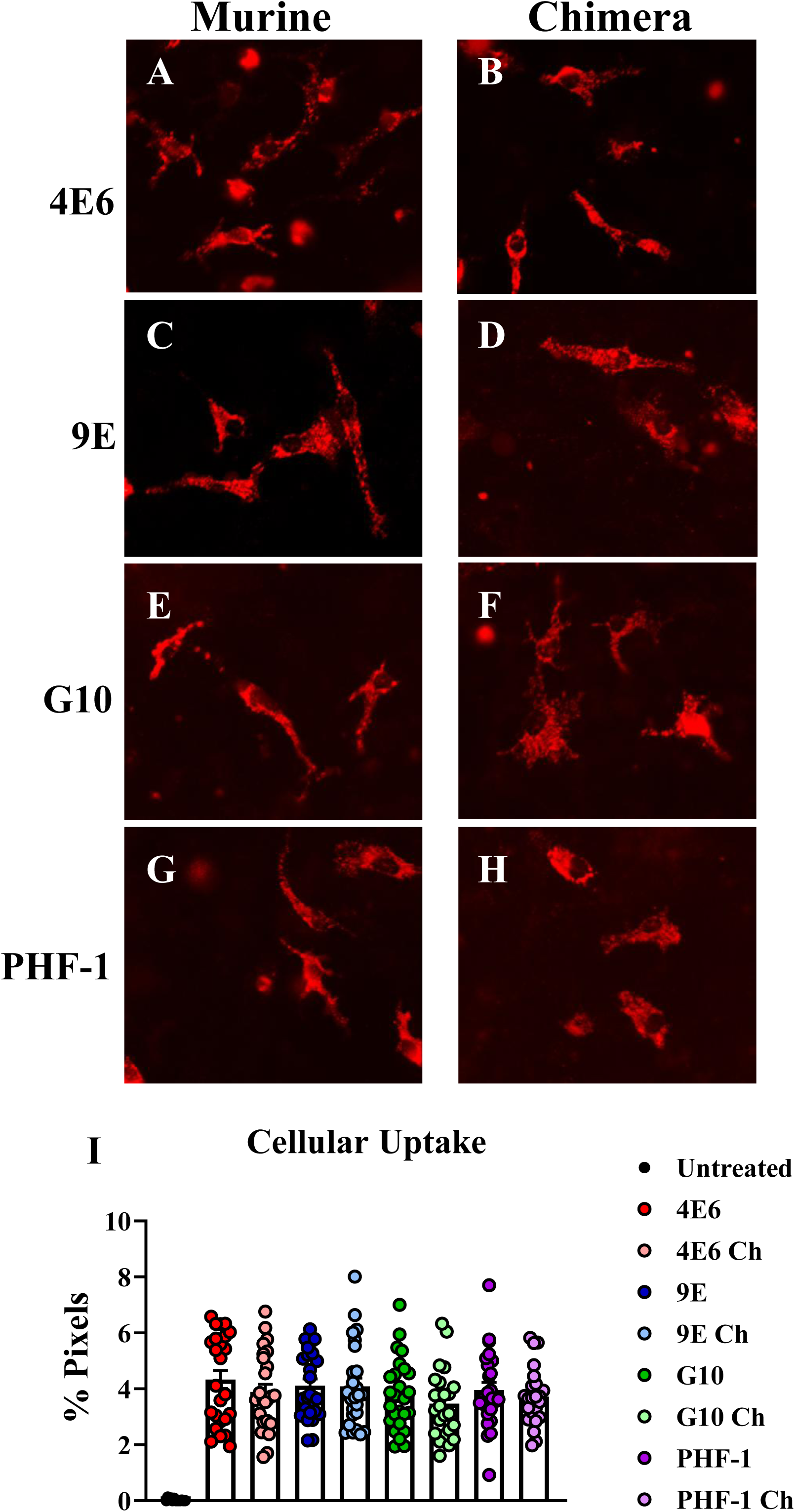
Murine and chimeric antibodies are internalized by cultured cells. Cortical cultures were incubated with 5 µg/ml CypHer5 labeled Abs for 1 hour, and then fixed with 4% PFA. **A-H**. Images were collected from each sample at 20X magnification. **I.** The percentage of pixels containing antibody fluorescence for each image was determined. All the Abs were taken up to a similar extent with no significant differences between the groups.

We then compared the efficacy of the two 4E6 chimeras. Mixed cortical cultures were prepared and treated using the PHF → Ab paradigm as described above. Cell lysate was probed for GAPDH and total tau (**Figure S4C**). Incubation with PHF alone resulted in reduced GAPDH levels in the cells (34% control values, p < 0.0001). Neither form of chimeric 4E6 prevented this toxicity (GAPDH 33 and 30% control values for F and S respectively, p < 0.0001 for both). Intracellular tau levels were increased following PHF exposure (Total tau/GAPDH ratio 2.76, p = 0.0008). As with GAPDH, neither chimeric 4E6 Ab was effective in preventing seeding (Total tau/GAPDH ratio 2.1 and 2.3, p = 0.01, = 0.005 for F and S respectively, **Figure S4D**).

These data suggest that even when the Ab is internalized, the unfavorable binding profile of chimeric 4E6 renders it ineffective in preventing PFH-induced toxicity. While it may show some efficacy in preventing the uptake of some non-toxic tau species in the PHF + Ab paradigm, as shown in previous results and **Figure 6**, it is unable to promote the clearance of internalized tau in the endosomal-lysosomal system. Thus, the differences between the uptake results is likely due to changes in the cell type used to generate the Ab. Regardless, neither 4E6 chimera prevented tau toxicity and seeding in the intracellular paradigm.

### Crystal structure of Fab PHF-1 in complex with a phosphorylated Tau epitope pS396

To understand the antigen-Ab interactions of 9E, G10, and PHF-1, we attempted to crystallize their Fab fragments in complex with their epitope peptides with either one or two phosphorylated sites. However, we found in our first atempt that all crystals we obtained were the Fab alone (**Table 4, Figures S5 and S6**). The antigen bindings sites revealed clear features associated with phospho-serine (pSer) recognition (**Figure S6**), including positively charged residues. Each binding site contains multiple positively charged residues: 9E and G10 both have two Arg and one Lys, while PHF-1 contains three Arg and two Lys residues. Although the CDR L3 and H3 loops are relatively short (**Supplemental Table 1**), the L1 and H1 loops are extended and form a crevice at the binding site, likely accommodating the peptide epitope (**Figures S5 and S6**). Notably, unlike mAb 4E6 [49], the antigen-binding sites of 9E, G10, and PHF-1 do not contain deep pockets. This suggests that their pSer epitopes likely lie across the surface of the binding site and traverse the crevice formed between the L1 and H1 loops.

**Table 4.** Data collection and refinement to statistics of the Fab structures.

| **Data collection** | **PHF-1 Fab/pSer396** | **PHF-1 Fab** | **G10 Fab** | **9E Fab** |
| --- | --- | --- | --- | --- |
| Space group | P 1 21 1 | P 2 21 21 | P 21 2 21 | I 41 2 2 |
| Cell dimensions |  |  |  |  |
| a, b, c (Å) | 91.003 120.505 142.399 | 40.31 92.59 137.60 | 85.64 105.29 132.11 | 152.19 152.19 117.80 |
| $\alpha, \beta, \gamma$ (°) | 90 106.86 90 | 90.00 90.00 90.00 | 90.00 90.00 90.00 | 90.00 90.00 90.00 |
| Resolution (Å) | 2.55 (2.70-2.55) | 1.87 (1.92-1.87) | 3.02 (3.20-3.02) | 2.83(2.90-2.83) |
| Rmerge | 19.0 (85.3) | 6.0 (61.0) | 15.4 (76.9) | 18.1(107.3) |
| I / σI | 7.65 (1.91) | 24.1 (3.4) | 12.90 (2.47) | 14.6 (2.6) |
| Completeness (%) | 99.6 (98.7) | 99.6 (95.6) | 98.9 (94.7) | 99.6 (97.7) |
| Redundancy | 4.7 (4.7) | 13.4 (13.1) | 6.4 (6.2) | 14.9 (13.5) |
| **Refinement** |  |  |  |  |
| Resolution (Å) | 35.40-2.55 | 29.68-1.87 | 29.57-3.02 | 29.47-2.83 |
| Number of reflections | 95,790 | 43,335 | 44,602 | 16,797 |
| Rwork / Rfree | 19.6/25.5 | 19.7/23.2 | 19.6/25.2 | 19.2/25.0 |
| Number of atoms: |  |  |  |  |
| macromolecules | 20411 | 3353 | 6,418 | 3344 |
| ligand |  | 0 | 0 | 24 |
| Solvent | 1115 | 289 | 0 | 33 |
| Average B-factor | 39.63 | 37.40 | 61.04 | 57.5 |
| macromolecules | 39.67 | 36.92 | 61.04 | 57.5 |
| ligands |  | 0 | 0 | 60.7 |
| solvent | 38.88 | 43.01 | 0 | 52.4 |
| R.m.s. deviations |  |  |  |  |
| Bond lengths (Å) | 0.0086 | 0.006 | 0.008 | 0.008 |
| Bond angles (°) | 1.070 | 0.88 | 1.188 | 0.99 |
| Molrep model | 4WEB | 4WEB | 7D9Z | 7D9Z |
| PDB ID | 9Z0S | 9Z0U | 9Z0T | 9Z0V |

Statistics in parentheses refer to outer resolution shell.

We were finally able to determine the co-crystal structure of Fab PHF-1 bound to a phosphorylated tau peptide encompassing residues 386–408 (sequence: TDHGAEIVYK(pSer)PVVSGDTSPRHL). The structure was refined to 2.55 Å resolution with R_work_ = 19.6% and R_free_ = 25.5% (**Table 4**). The crystals belong to space group P21 and contain six Fab:peptide complexes in the asymmetric unit. Although the crystallization peptide was 23 residues long, only about nine residues (tau 391–399; sequence EIVYKpSPVV) were clearly resolved in the electron-density map of the 6 complexes in the asymmetric unit (**Figure 9A**). The electron density quality varies slightly among the complexes, which are otherwise identical, with an average root-mean-square deviation (RMSD) of 0.2 Å. For clarity, only one representative complex is described below. Ab residue numbering follows the Kabat convention, with heavy and light chains shown in green and cyan, respectively, throughout the figures. The light chain and heavy chain are labeled as H or L, respectively.

**Figure 9.**
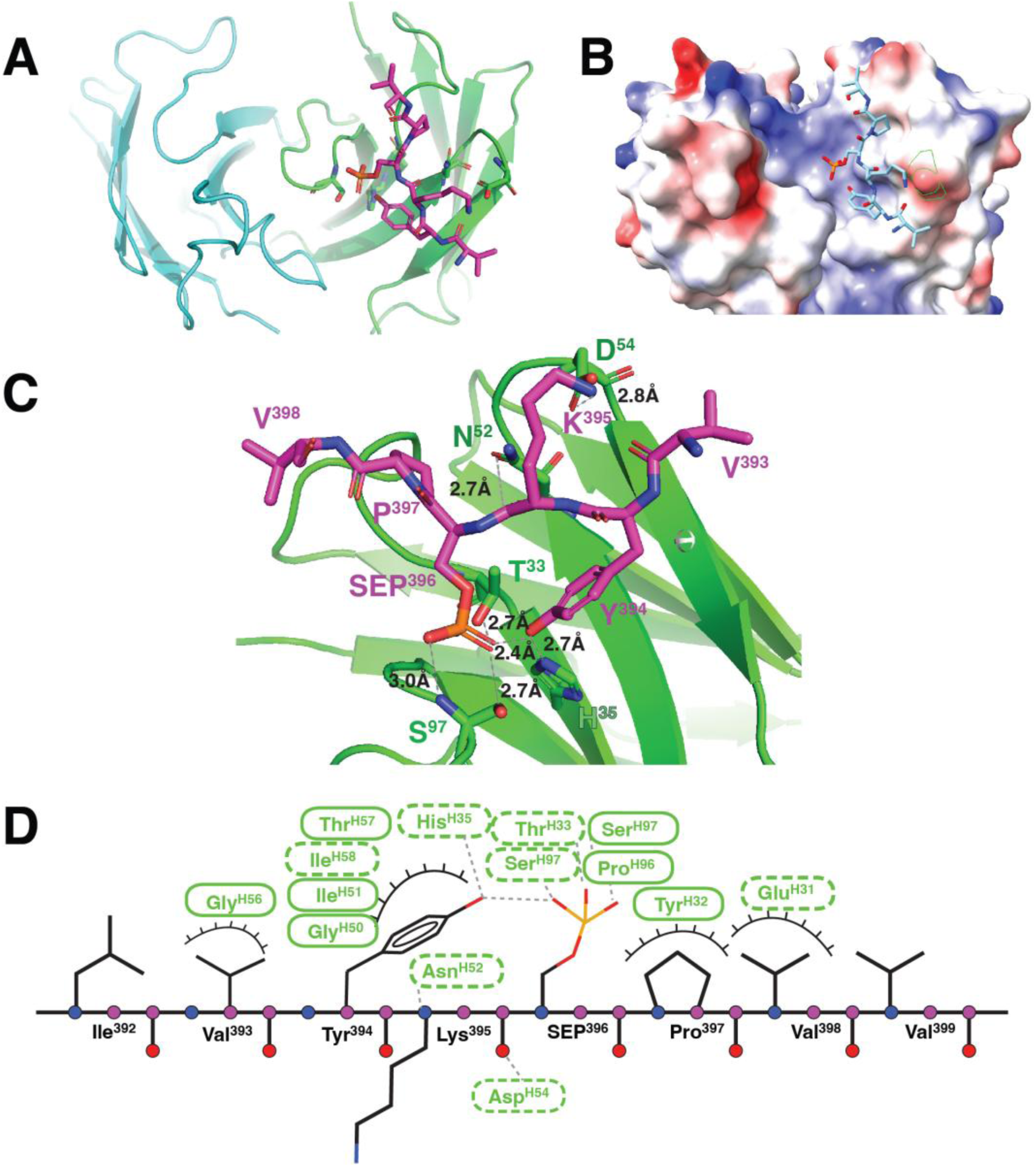
Structure of Fab PHF-1 in complex with pSer396 Tau peptide. **A**. Front-view ribbon representation of the Fab PHF-1:pSer396 complex. Although the crystallization peptide contained 23 residues, only residues EIVYKpSPVP (391–399) were resolved (shown as magenta sticks). Heavy and light chains are colored light green and cyan, respectively; note the peptide’s minimal contact with the light chain. **B**. Electrostatic surface of the PHF-1 antigen-binding site, colored from negative (red) to positive (blue). The strong electropositive region corresponds to the phosphate-binding pocket. **C**. Details of the antibody-antigen interactions. Hydrogen bonds are shown as dashed lines and labeled with bond distances. **D**. Schematic diagram of PHF-1 antibody–antigen interactions. Hydrogen bonds are shown as dashed lines; van der Waals contacts are indicated by eyelashes. Solid ovals represent main-chain interactions, while dashed ovals represent side-chain interactions.

PHF-1 has a heavy-chain-dominant binding mode (**Figure 9A**). The light chain contributes minimal contact to the tau peptide, whereas two tau residues, Tyr394 and pSer396, are deeply buried within a pocket formed by the heavy-chain complementarity-determining regions (CDRs). The Ab-antigen interface is stabilized by a network of hydrogen bonds, salt bridges, and hydrophobic interactions. The phosphorylated pSer396 residue is accommodated in an electropositive phosphate-binding pocket formed primarily by heavy-chain residues (**Figure 9B**). Residues from CDR-H1 and CDR-H3 confer phospho-specific recognition of pSer396. The phosphate group is anchored by three hydrogen bonds involving ThrH33 (CDR-H1) and SerH97 (CDR-H3). Specifically, the side chains of ThrH33 and SerH97 form direct hydrogen bonds with the phosphate oxygens of pSer396, while the backbone amide of SerH97 donates an additional hydrogen bond to the phosphate (**Figure 9C**). A salt bridge is formed between AspH54 and Lys395, and AsnH52 donates a hydrogen bond to the main chain of Lys395. HisH35 recognizes the aromatic ring of Tyr394 through polar and π-stacking interactions.

Within the complex, the peptide backbone adopts a conformation stabilized by a Tyr394-pSer396 side-chain hydrogen bond, effectively locking the local structure. The orientation of Tyr394 differs from that in unbound tau, flipping to enable interaction with SerP396. This observation supports our previous finding [36] that the YXSP motif in tau acts as a phosphorylation-dependent conformational switch, converting local structure from a potentially α-helical to β-strand state upon serine phosphorylation. The present structure thus provides a mechanistic explanation for PHF-1’s specificity toward pathological, hyperphosphorylated tau.

## Discussion

Using DNA prime-protein boost regimen, which initiates antigen expression in vivo and is followed by peptide boosting to strengthen antibody responses [77], mice were immunized with a DNA plasmid encoding the 2N4R full length tau isoform. The animals were then boosted using peptides containing tau residues 391-407 (EIVYKpSPVVSGDTpSPRH) conjugated to an HIV-1 gp120 T1 peptide (KQIINMWQEVGKAMYA) at the N-terminal, or tau residues 394-406 conjugated to a tetanus toxin epitope region (GPSLFNNFTVSFWLRVPKVSASHLE) [56]. From this approach we generated two new mAbs 9E and G10 targeting pSer404 and pSer396, respectively. We compared their epitope specificity, efficacy in culture, and tau binding along with 4E6, PHF-1, and their partially humanized chimeric variants. 4E6 and 9E showed high affinity for the pSer404 peptide, and not the pSer396 peptide, while the converse was true for G10 and PHF- 1. The pattern of peptide binding was the same for the chimeric variants. Additionally, we crystalized the Fab regions of each Ab to compare the binding surface at an atomic level. Similar to our previously published results with 4E6 [48], chimerization increased the isoelectric point of all the Abs. Interestingly, the impact of chimerization on Ab efficacy against PHF-induced toxicity and seeding, as well as tau binding differed substantially.

Consistent with our previous results, murine 4E6 was effective in both extra- (PHF + Ab) and intracellular (PHF → Ab) dosing paradigms, with robust cellular uptake and preferential binding to soluble tau aggregates, whereas its chimera blocked tau seeding but did not prevent tau toxicity in the PHF + Ab dosing method [48]. Murine 9E prevented tau toxicity and seeding in both the PHF + Ab and PHF → Ab dosing paradigms, while its chimera prevented toxicity and seeding only in the intracellular PHF → Ab dosing method. Chimeric G10 also prevented PHF-induced loss of GAPDH and increase of tau in both the extra- and intracellular paradigms, but the murine G10 only prevented tau seeding in the PHF + Ab treatment. Murine PHF-1 prevented toxicity and seeding extracellularly, while the chimeric variant did not prevent toxicity or seeding by either dosing method. Overall for efficacy, the chimerization was detrimental for 4E6 and PHF-1, beneficial for G10, and neutral for 9E. These results also demonstrate that prevention of tau toxicity and seeding do not necessarily go hand in hand and that both should be assessed when determining Ab efficacy.

We additionally tested two other mAbs, 6B2 and 8B2. In previous experiments 6B2 bound non-phosphorylated, pSer396/pSer404, pSer396, and pSer404 to a similar extent in solid phase ELISA, and showed only partial phospho-selectivity in a competitive solution phase ELISA [44]. We have published results showing phospho-selectivity of 8B2 [35]. In solid phase ELISA 8B2 bound strongly to non-phosphorylated, pSer396/pSer404, and pSer404 peptides with binding to pSer396 only at high Ab concentrations, and in the solution phase binding was inhibited to a much greater extent using a pSer/396/404 peptide compared to the non-phosphorylated peptide [35]. In this study we examined 6B2 and 8B2 binding to pathological tau using a competitive ELISA. Binding to the plate was inhibited when 8B2 was preincubated with solubilized PHF, but there was no effect on 6B2 binding. When we examined the efficacy of the two Abs, 6B2 was unable to prevent tau-induced toxicity and seeding in either dosing paradigm, consistent with our previous findings [49]. In contrast, 8B2 prevented toxicity and seeding in the PHF + Ab paradigm and seeding in the PHF → Ab method. This again highlights that high affinity for the antigen does not necessarily translate into efficacy, and that preferential binding to soluble tau species is a more reliable marker.

Interestingly, we found that the cellular uptake data differed from that seen previously. We found that all Abs displayed a similar level of uptake. While we did not examine the specific cell types in this experiment, in previous studies we have found that in that uptake is primarily neuronal. In cultures containing neurons and glia and no exogenous PHF, approximately 80% of Ab uptake is neuronal, followed by 10% astrocytic, and the remaining 10% other cell types, and that this is important for efficacy [44, 48]. In past experiments chimeric 4E6 showed only limited cellular uptake while in this study it was comparable to its murine form. We hypothesize that this may have been linked to the cell type used to generate the Ab, as both 293F and 293S cells were being used at the time [35, 48]. The 293S cells have a simplified glycosylation pattern relative to the 293F cells [58]. This is desirable when generating proteins for crystallization, but changes in glycosylation can impact Ab-receptor binding. While we confirmed that the previous efficacy testing had been conducted using Ab made in 293F cells, we did not have any of the prior CypHer5 labeled chimeric 4E6 Ab to examine. We generated new chimeric 4E6 Abs from both 293F and 293S cells and tested them for uptake and binding. Uptake was reduced in the cultures incubated with chimeric 4E6 from 293S cells compared to that from 293F cells, suggesting that the difference from previous results may be due to the use of Abs from two different cell lines. Neither prevented toxicity or seeding induced by intracellular PHF, indicating that even if the chimeric 4E6 is internalized it is still ineffective. In the extracellular binding paradigm, chimeric 4E6 also failed to prevent toxicity and its binding profile showed preferential binding to insoluble tau species (**Figure 3A, 4A**, [48]).While it prevented the uptake of some non-toxic tau species in the extracellular paradigm it was not able to facilitate clearance of already internalized PHF. Like 6B2, its overall higher tau binding may actually prevent the breakup of aggregates or block access to lysosomal enzymes. Neuronal uptake is still an important feature, as we have shown that blocking uptake of murine 4E6 reduces its efficacy in the PHF → Ab paradigm [48]. However, here the efficacy differences between murine and chimeric 4E6 can be explained by differences in their tau binding profile.

The competitive ELISA data provided further insight into why some Abs were more effective. In past experiments, Abs with decreased solid phase binding after pre-incubation with PHF-tau, such as murine 4E6, were efficacious, whereas those that maintained the solid phase binding, 6B2 and chimeric 4E6, were ineffective [47, 48]. Small soluble oligomers are the more toxic tau species, thus Abs that bind to them are better at preventing tau-induced toxicity. Murine and chimeric 9E both had reduced solid phase binding after pre-incubation with PHF-tau, and both were efficacious in the PHF → Ab paradigm. Similarly, chimeric G10 and murine PHF-1 had decreased solid phase binding to a greater extent than their other species pair. This may explain why murine G10 did not prevent tau toxicity in either dosing method, or why murine but not chimeric PHF-1 prevented toxicity in the PHF + Ab paradigm. With 9E, it is possible that there are some additional subtle changes in binding that impaired its efficacy in the PHF + Ab paradigm. In the extracellular space it may bind to a non-toxic species, but when it is internalized and in the same compartment as the pathological tau, it can bind to and clear or neutralize it.

The crystal structure of Fab PHF-1, a widely used antibody in the field, bound to the tau pSer396 epitope (EIVYKpSPVV; 2.55 Å resolution) shows that the peptide lies across the top of the antigen-binding surface, with minimal contact from the light chain, confirming PHF-1 as a heavy-chain-dominant Ab. The pSer396 residue is anchored in an electropositive pocket via several hydrogen bonds to the phosphate moiety, conferring phospho-specificity. Adjacent Lys395 and Tyr394 contribute additional stabilization via salt-bridge and π-stacking interactions with AspH54, AsnH52, and HisH35. Notably, TyrP394 flips its side-chain orientation to hydrogen-bond with pSer396, locking the local backbone into a β-strand-like conformation. This structural rearrangement corroborates our earlier biochemical finding that the YXSP motif functions as a phosphorylation-dependent conformational switch, converting α-helical segments of tau into β-strand structures characteristic of pathological aggregates. This unique binding pattern may explain the efficacy profile seen with this Ab. Murine PHF-1 had a favorable binding profile in the ELISAs and was effective in the extracellular PHF + Ab paradigm. Despite this, it could not prevent toxicity when the PHF was already internalized. If the Ab promoted misfolding or stabilized already misfolded tau, this may explain why. PHF-1 is capable of blocking the uptake of toxic tau into neurons but has limited ability to facilitate removal of already internalized tau species.

Our structural data unify the electrostatic and conformational principles underlying pSer recognition among this Ab family. While the unliganded structures of 9E and G10 predicted a shallow, charge-complementary mode of recognition, the PHF-1:pSer396 complex now visually confirms this model at atomic resolution. The combination of a positively charged surface, short CDR loops, and a planar binding groove emerges as a recurring architectural solution for Abs targeting phosphorylated tau epitopes-balancing specificity for the phosphate group with adaptability to local conformational changes within tau.

Together these data raise important points regarding the development of tau Abs as therapeutics. For example, while chimeric 4E6 and murine G10 prevented intracellular tau seeding, they did not ameliorate the toxic effects of PHF, which supports our long argued point that efficacy should not be measured by inhibition of tau seeding alone.

Epitope selection is a crucial aspect of Ab design, and the choice of target can dictate whether the Ab is effective or not. While it may initially appear that targeting pSer404 is more efficacious based on the results using 4E6 and 9E, the interpretation requires more nuance. While murine G10 was ineffective, chimeric G10 prevented toxicity and seeding in both dosing paradigms. As discussed above, the difference is likely due to changes in tau binding following chimerization. Thus, various properties of individual Abs determine their efficacy. Overall, these data, along with our previous work and that of others, show the validity of targeting the pSer396/pSer404 epitope and that targeting either specific site can prevent tau seeding, [17, 39–41, 43, 46, 48–50]. The data presented herein and previously published reports also show that targeting both epitopes can also prevent toxicity [48, 49], although targeting pSer404 appears to be generally more efficacious than targeting pSer396 based on our current findings. We would again emphasize the need to consider both toxicity and seeding when determining antibody efficacy as the two measures may not always be in concert.

Finally, a point we have also raised before is that chimerization should also not be treated as a simple process of switching IgG. As seen here, and in our previous results, changing IgG species or subclass can change tau binding and efficacy even when the Fab region is identical [48, 52]. This is supported by research from other groups showing that changing the Fc region of the Ab can impact target engagement despite unchanged variable regions [64–66, 69–76, 78–82]. All parts of the Ab contribute to binding, uptake, and clearance, and should not be treated as discrete entities that can be swapped indiscriminately. This has been more thoroughly explored in the cancer immunotherapy field, and the lessons from those findings should be applied to the development of tau targeting Abs as well.

## Acknowledgement

We thank Peter Davies (Albert Einstein College of Medicine and Long Island Jewish Medical Center, Litwin-Zucker Research Center at Feinstein Institutes for Medical Research) for providing the purified PHF-1 tau antibody. We thank the NDRI (Philadelphia, PA), from which we purchased human brain tissue samples collected from a patient with mixed Alzheimer’s disease/Pick’s disease pathology. This research used resources beam line 17–ID-1 (AMX) of the National Synchrotron Light Source II, a U.S. Department of Energy (DOE) Office of Science User Facility operated for the DOE Office of Science by Brookhaven National Laboratory under Contract No. DE-SC0012704. The Center for BioMolecular Structure (CBMS) is primarily supported by the National Institutes of Health, National Institute of General Medical Sciences (NIGMS) through a Center Core P30 Grant (P30GM133893), and by the DOE Office of Biological and Environmental Research (KP1605010).

