## Supplemental Materials for "Tau pSer396 and pSer404 Define Distinct Epitope Regions Linked to Different Antibody Functions"

**Supplemental Table 1. Gene usages and CDR sequences (Kabat defininition) of 9E, G10, and PHF-1.**

| Heavy chain | Gene Usage | CDRH3 |
|---|---|---|
| 9E | IGHV7-3*02 | DRGLTFGYDWFAY |
| G10 | IGHV5-6*01 | HNDFHAMDY |
| PHF-1 | IGHV1-18*04 | GPSARFPY |

| Light chain | Gene Usage | CDRL3 |
|---|---|---|
| 9E | IGKV1-133*01 | VQGTHFPYT |
| G10 | IGKV1-117*01 | FQASHVPWT |
| PHF-1 | IGKV1-135*01 | WQGTHFPRT |

**Supplemental Figure 1. 8B2 but not 6B2 showed decreased binding in a competition ELISA.** 6B2 and 8B were preincubated with increasing concentrations of PHF before being added to the coated ELISA plate. 6B2 showed no reduction in binding at any of the preincubation concentrations, while 8B2 had reduced binding at all but the lowest PHF dose (37-50% reduction, $p = 0.001$ - $< 0.0001$)
** $p \leq 0.01$, *** $p \leq 0.001$, **** $p < 0.0001$

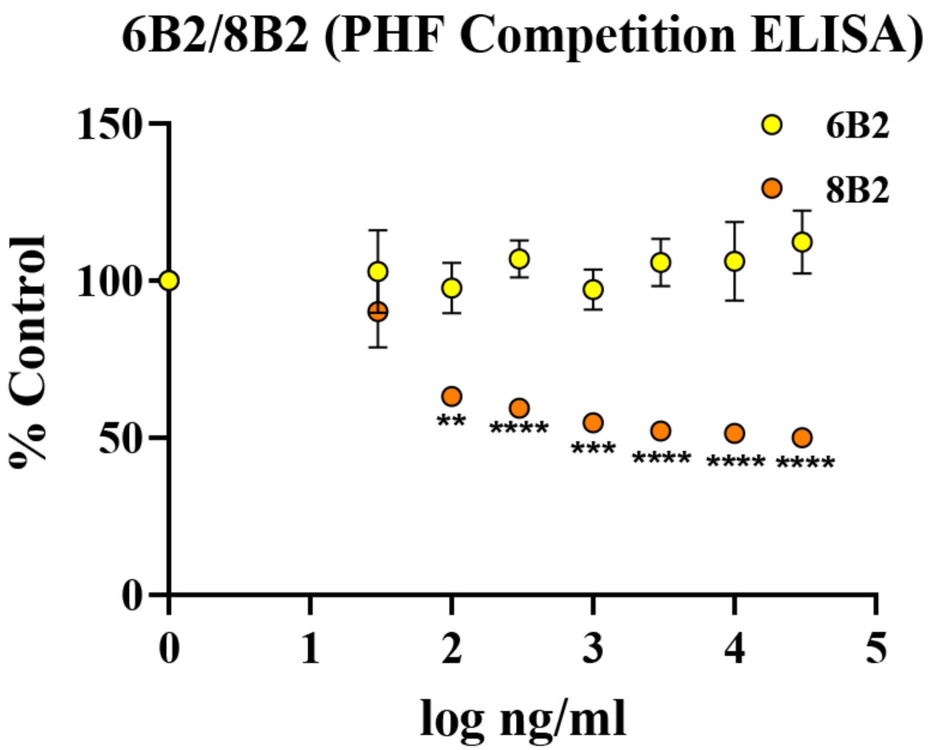

**Supplemental Figure 2. 8B2, but not 6B2 prevents PHF-induced toxicity and seeding in the PHF + Ab dosing paradigm, and seeding in the PHF → Ab dosing paradigm**. Mixed cortical cultures were treated with 10 μg/ml PHF and 10 μg/ml of either 6B2 or 8B2 using the PHF + Ab and PHF → Ab dosing paradigms. Samples were collected and assayed for levels of GAPDH and total tau. **A**. Immunoblots showing GAPDH and tau in samples from the PHF + Ab method. **B**. A significant overall treatment effect was seen by one-way ANOVA ($p < 0.0001$). PHF alone decreased GAPDH relative to untreated controls (41.4% of control values, $p < 0.0001$). GAPDH in samples treated with PHF and 6B2 was also reduced compared to control (56.7% of control values, $p = 0.0003$). When added to the cultures together with PHF, 8B2 prevented tau toxicity (141.2% of control values, $p < 0.0001$). **C**. There was a significant treatment effect on total tau levels ($p < 0.0001$). Cells treated with PHF alone and PHF in combination with 6B2 had significantly higher total tau levels compared to untreated controls (Tau/GAPDH ratio 2.0, 1.5, $p = 0.0002, 0.009$). 8B2 prevented this increase, resulting in lower total tau compared to PHF alone (Tau/GAPDH ratio 0.59, $p < 0.0001$). **D**. Immunoblots for GAPDH and total tau from cultures treated using the PHF → Ab dosing method. **E**.A one-way ANOVA showed a significant treatment effect ($p = 0.004$). Samples treated with both PHF alone and PHF followed by 6B2 had lower GAPDH levels compared to untreated controls ($p = 0.04, 0.003$). **F.** As with the other markers, a significant treatment effect was observed ($p = 0.003$). Cells exposed to PHF alone, and PHF followed by 6B2 had higher total tau (Tau/GAPDH ratio 2.4, 2.2, $p = 0.05$ for both). 8B2 treated samples had lower total tau compared to PHF alone (Tau/GAPDH ratio 0.59, $p = 0.02$).
### $p \leq 0.05$, ## $p \leq 0.01$, ### $p \leq 0.001$, #### $p < 0.0001$
* $p \leq 0.05$, **** $p < 0.0001$

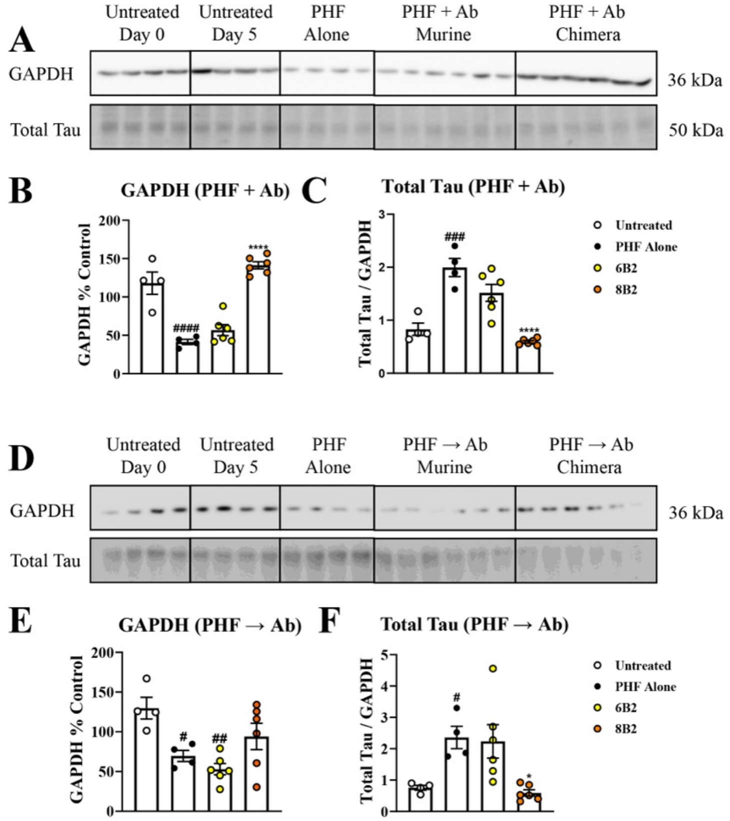

**Supplemental Figure 3. Endo Hf digestion of 4E6 chimera produced in 293S and 293F cells under denaturing conditions.** 4E6 chimera expressed in 293S GnTI-/- or 293F cells was analyzed by SDS-PAGE, including treatment with Endo Hf under denaturing conditions. Lanes 1–2, 4E6 chimera produced in 293S GnTI-/- cells without (lane 1) or with (lane 2) Endo Hf; lanes 3–4, 4E6 chimera produced in 293F cells without (lane 3) or with (lane 4) Endo Hf; lanes 5–6, mock protein (1FD6 scaffolded V1V2 of HIV-1 gp120) from 293S cells without (lane 5) or with (lane 6) Endo Hf. No clear difference in heavy-chain migration was observed between 4E6 chimera produced in 293S GnTI-/- and 293F cells under these denaturing SDS-PAGE conditions.

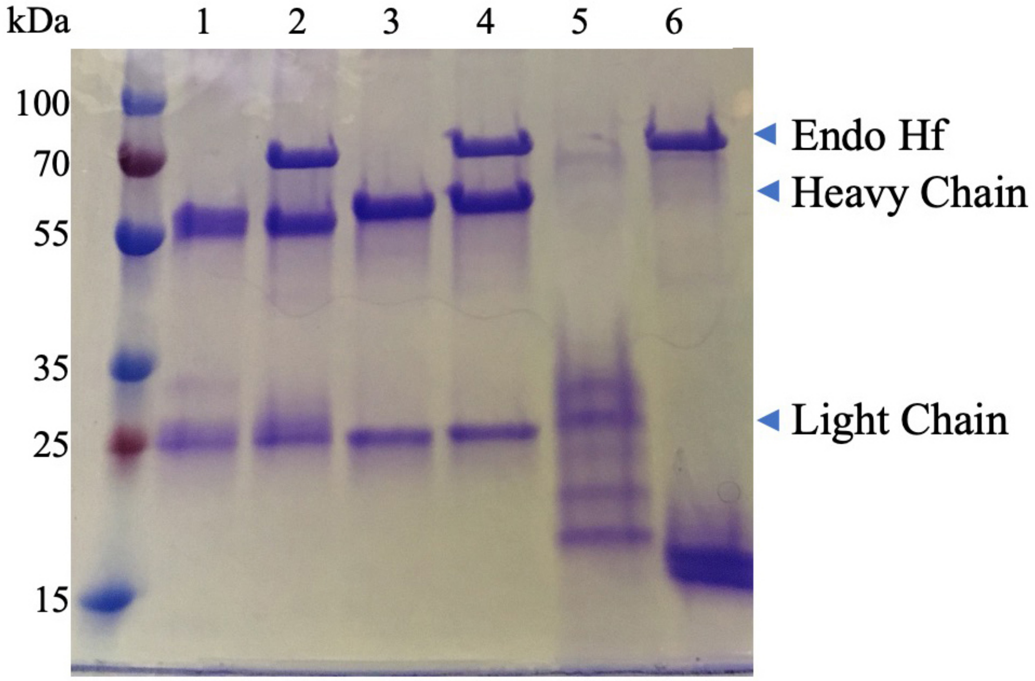

**Supplemental Figure 4. Chimeric 4E6 produced in 293S cells has reduced uptake compared to that produced in 293F cells. Neither Ab prevents tau-induced toxicity and seeding in the PHF → Ab paradigm. A**. Cortical cultures were incubated with 1 or 2 µg/ml CypHer5 labeled Abs for 1 hour and then fixed in 4% PFA. Images were collected from each sample at 20X magnification. **B.** The percentage of pixels containing antibody fluorescence for each image was determined. Cellular uptake was lower in cultures treated with 4E6 Ch S ($p < 0.0001$). **C**. Immunoblots showing GAPDH and total tau levels in cultures treated with PHF, 4E6 Ch F and 4E6 Ch S in the PHF → Ab dosing paradigm. **D**. Exposure to PHF alone reduced GAPDH levels (34.2 % of control values, $p < 0.0001$). Neither 4E6 Ch F or 4E6 Ch S prevented this toxicity (32.7 and 30.1 % of control values, $p < 0.0001$ for both). **E**. Intracellular tau levels increased following PHF exposure (Total tau/GAPDH ratio 2.76, $p = 0.0008$). Neither 4E6 Ab was effective in preventing seeding (Total tau/GAPDH ratio 2.1 and 2.3, $p = 0.01$, $= 0.005$ for 4E6 Ch F and S, respectively).

### $p \leq 0.05$, ## $p \leq 0.01$, ### $p \leq 0.001$, #### $p < 0.0001$

**** $p < 0.0001$

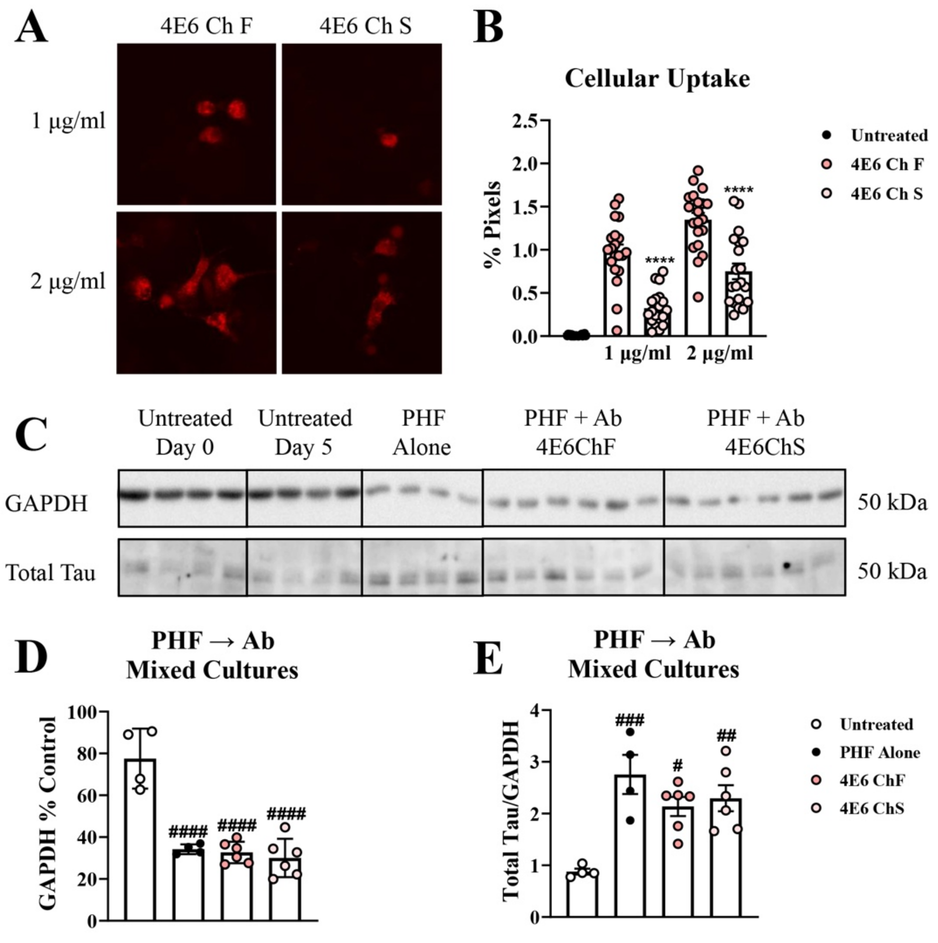

**Supplemental Figure 5. Ribbon representations of the crystal structures of Fabs 9E, G10, and PHF-1. A–C**. Front and side views of Fabs 9E (**A**), G10 (**B**), and PHF-1 (**C**), respectively. Light chains are shown in cyan and heavy chains in green.

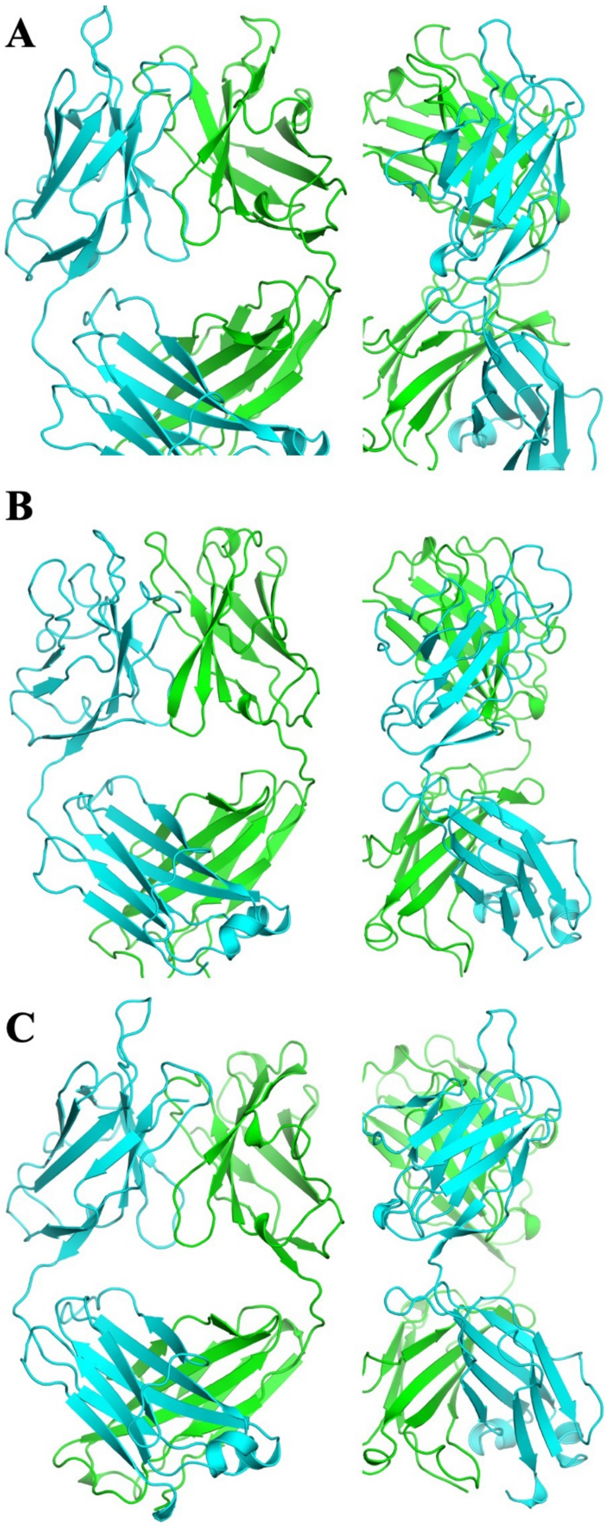

**Supplemental Figure 6. Structural features of the antigen-binding sites of mAbs 9E, G10, and PHF-1. A**. Top views of the antigen-binding sites of the three mAbs shown in ribbon representation. The three CDR loops of the light chain are colored in marine, light blue, and deep blue, while those of the heavy chain are shown in light orange, orange, and olive, respectively. **B.** Electrostatic potential surfaces of the antigen-binding sites, highlighting the overall positively charged nature of these surfaces.

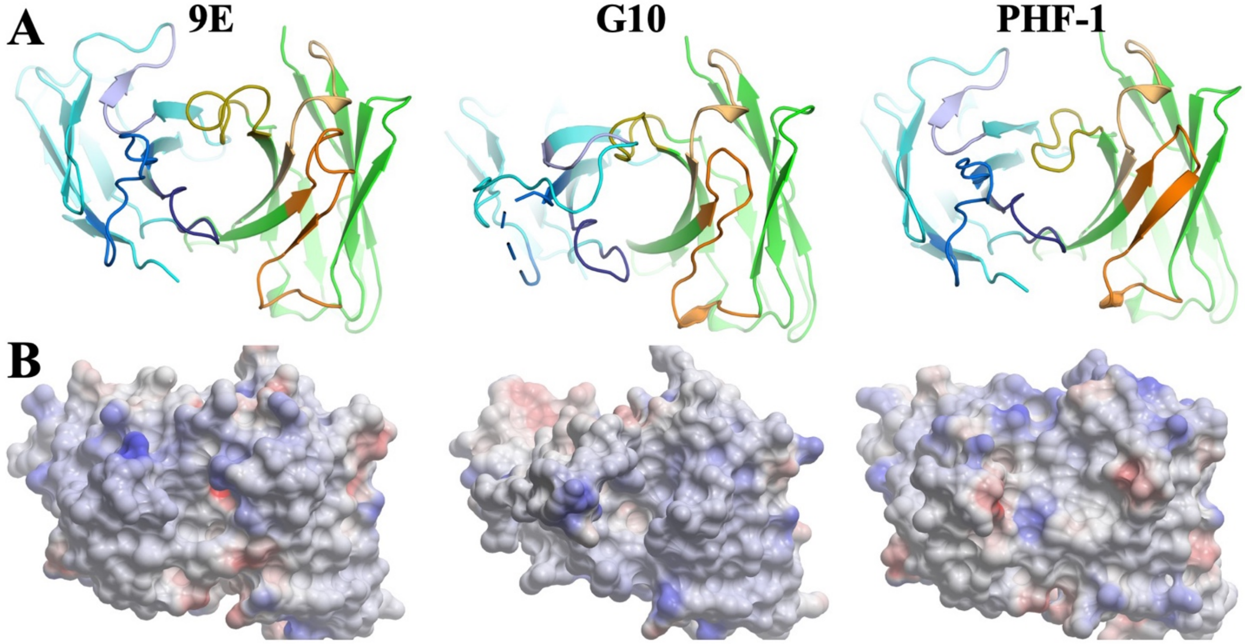
